# Lithocholic acid induces T3SS-dependent formation of invasion-competent *Shigella flexneri* aggregates

**DOI:** 10.1101/2025.11.24.690263

**Authors:** Jonah Lanier, Jaden J. Skelly, Freddie Salsbury, Volkan K. Köseoğlu

## Abstract

*Shigella flexneri* causes shigellosis, the second leading cause of diarrheal deaths worldwide. The pathogen invades colonic epithelial cells using a type III secretion system (T3SS) that delivers effector proteins to remodel host actin cytoskeleton. Following invasion, *S. flexneri* acquires actin-based motility and spreads cell to cell, driving epithelial destruction and bloody diarrhea. These intracellular infection processes have been investigated primarily using exponentially growing planktonic bacteria. However, recent animal studies revealed that *S. flexneri* also forms multicellular aggregates in the colonic lumen, yet the function of this extracellular phase remains unclear. Here, we show that lithocholic acid (LCA), an abundant secondary bile acid in the colon, acts as a potent signal that induces *S. flexneri* aggregation at physiological concentrations (≥50 µM). LCA-induced aggregation depends on the T3SS and its tip protein IpaD, which increases aggregate size. Compared to non-aggregating controls, LCA-induced aggregates initiate invasion by eliciting a more robust actin remodeling and rapid T3SS activation during early interactions with colonic epithelial HT-29 cells. These findings identify LCA as a luminal cue that links the extracellular aggregation with intracellular infection, through a new aggregate-mediated mode of epithelial invasion.

## INTRODUCTION

*Shigella* spp. are the causative agents of bacillary dysentery, or shigellosis, a highly contagious diarrheal disease characterized by abdominal cramps, fever, severe diarrhea, and stools containing mucus and blood (1,2). Globally, shigellosis accounts for an estimated 270 million cases and 200,000 deaths annually, disproportionately affecting children under five, with 64,000 deaths and long-term consequences such as persistent diarrhea and growth faltering (3–7). Despite ongoing vaccine development efforts, no licensed vaccine is currently available (8, 9). The World Health Organization recently designated *Shigella* spp. as a “high priority” pathogen due to rising antimicrobial resistance (10).

Among *Shigella* spp., *S. flexneri* is the predominant cause of shigellosis, responsible for 65% of pediatric and 60% of all cases worldwide (6, 11). It is highly infectious, with as few as 100 bacteria sufficient to cause disease (9). Transmission occurs through person-to-person contact, ingestion of contaminated food or water, and, in some cases, sexual contact (1). Following transmission, *S. flexneri* ultimately invades the colonic epithelium, where it causes mucosal ulceration marked by inflammation, vascular damage, and epithelial detachment, hallmarks confirmed in human biopsy studies (12–15). Invasion of epithelial cells relies on type III secretion system (T3SS) of the pathogen, which delivers effector proteins into host cells to subvert cellular processes (16). Once inside the cytosol, *S. flexneri* uses the polar surface protein IcsA to acquire actin-based motility, enabling intracellular movement (17, 18). Actin-based motility, combined with effector-driven manipulation of plasma membrane, enables the pathogen to disseminate across the colonic epithelium (19, 20).

While the intracellular phase of *S. flexneri* infection has been extensively studied, mechanisms supporting the extracellular phase remain poorly understood. The low infectious dose and an incubation period of up to four days (1, 21) suggest that *S. flexneri* establishes extracellular colonization in the colon. Supporting this, recent studies in infant rabbit and mouse models have shown that *S. flexneri* replicates extracellularly and forms dense aggregates in the colonic lumen (22, 23). One class of luminal molecules that may influence this early colonization phase is bile acids (24). These detergent-like compounds are synthesized from cholesterol in the liver as primary bile acids, stored in the gallbladder, and secreted into the small intestine within bile following food intake (25). There, they facilitate dietary lipid digestion and absorption (26). Approximately 95% of bile acids undergo enterohepatic circulation, while the 5% remains in the intestines, where gut microbiota convert primary bile acids into secondary bile acids (27). As a result, bile acid composition shifts dramatically between the small intestine and the colon. Primary bile acids, cholic acid and chenodeoxycholic acid, dominate in the small intestine, whereas secondary bile acids, deoxycholic acid (DCA) and lithocholic acid (LCA), are more abundant in the colon (28). Beyond their role in lipid digestion, bile acids act as signaling and antimicrobial compounds, shaping gut microbiota composition and reinforcing colonization resistance (29–32). Pathogens like *S. flexneri* have evolved strategies to resist bile acid stress, including efflux pumps, membrane remodeling, and biofilm formation (24). Previous studies have shown that DCA promotes *S. flexneri* aggregation and biofilm formation (33–35), modulates virulence factor expression and epithelial adherence (36–38). However, the role of LCA, another predominant colonic bile acid, remains unexplored. In this study, we investigated how LCA influences *S. flexneri* during extracellular growth and its early interactions with epithelial cells. We demonstrate that LCA triggers T3SS-dependent aggregation and that these aggregates retain the capacity to invade epithelial cells. Together, these findings identify LCA as a luminal cue that drives the formation of invasion-competent aggregates, suggesting a potential role for aggregates at the interface between extracellular colonization and epithelial invasion.

## RESULTS

### Lithocholic acid promotes aggregation in *S. flexneri*

Deoxycholic acid (DCA) and lithocholic acid (LCA) are the two predominant secondary bile acids in the colon (28). While DCA has been previously shown to promote biofilm formation and aggregation under static growth conditions (33–35), the impact of LCA on *S. flexneri* community behavior had not been examined. We first assessed biofilm formation in Tryptic Soy Broth supplemented with increasing concentrations of LCA (10–2500 μM) using crystal violet staining. Across this range, LCA had no measurable effect on biofilm development on polystyrene surfaces (**Fig. 1A**), in contrast to DCA (**Fig. 1B**). We next tested the effect of LCA on *S. flexneri* aggregation. LCA induced readily visible aggregation at concentrations as low as 50 μM, forming clumps at the bottom of culture tubes after 24 hours of static incubation at 37°C (**Fig. 1C**). However, DCA induced aggregation only at substantially higher concentrations (500-2500 μM) (**Fig. 1D**), indicating that LCA is drastically more potent at inducing aggregation. To validate OD_600_ as a measure of aggregation, we independently quantified aggregation using CFUs. Both methods yielded similar planktonic-to-aggregate ratios (**Fig. S1**), confirming OD_600_ as a reliable readout.

**Figure 1.**
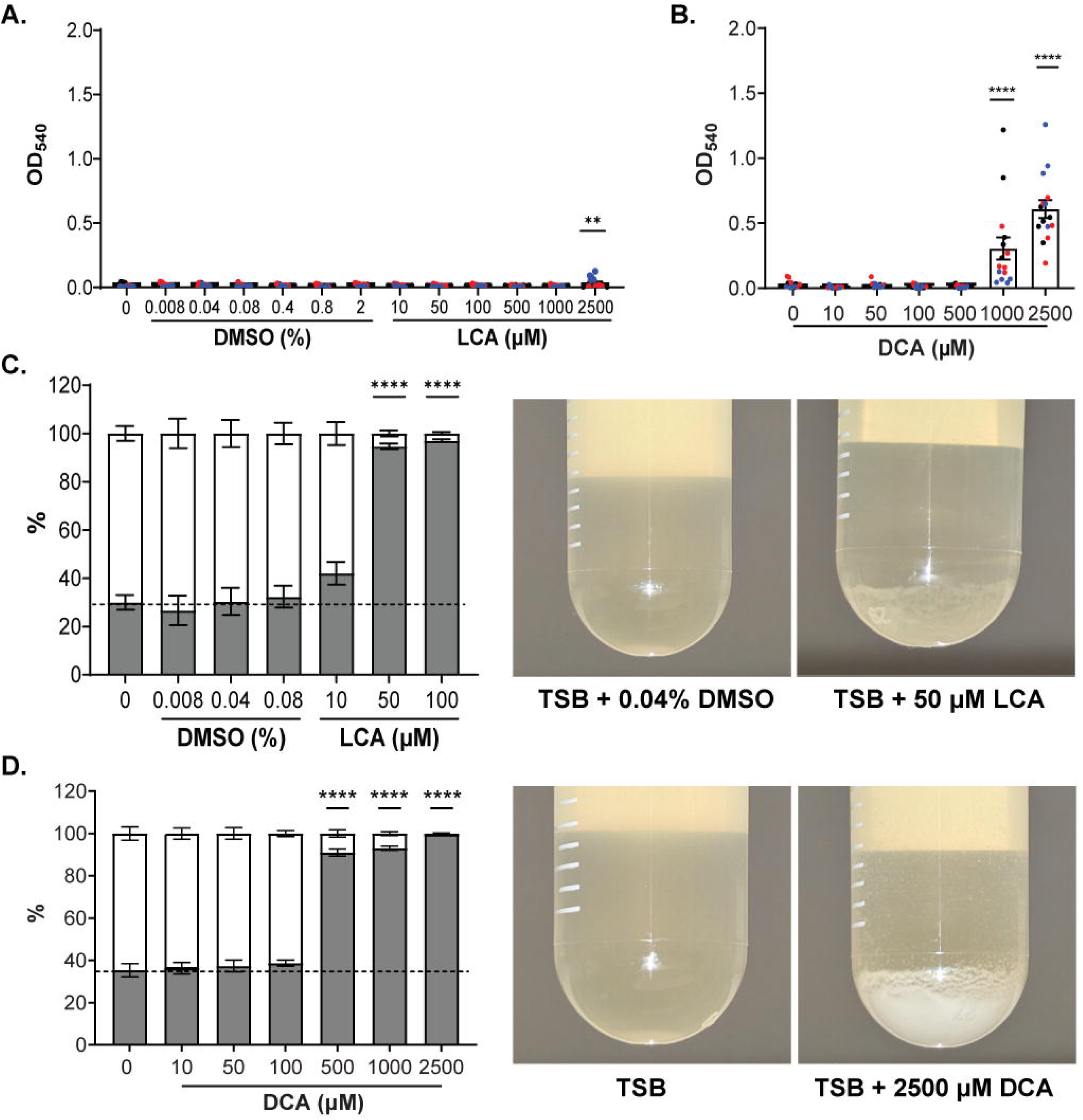
LCA induces robust aggregation in standing *S. flexneri* cultures. (A) Crystal violet staining of biofilm after 24 h growth across increasing LCA concentrations, each paired with its matched DMSO control. B) Biofilm biomass quantified across increasing DCA concentrations. Graphs show mean ± SEM; each color represents an independent biological repeat (n =3). (C) Sedimentation assay of S. *flexneri* in TSB containing increasing concentrations of LCA and vehicle (DMSO). (D) Sedimentation assay of S. *flexneri* grown in TSB with increasing concentrations of DCA. Graphs show mean ± SD from three biological repeats; white bar, planktonic %; gray bar, aggregated %. Dashed line indicates basal bile acid independent aggregation. Representative culture tubes are shown for comparison. Statistics: one-way ANOVA with Dunnet’s multiple comparison test. **, p<0.01; ****, p<0.0001. Each LCA concentration was compared to corresponding DMSO %. Each DCA concentration was compared to the TSB control.

Because bile acids can exhibit antimicrobial activity, we next asked whether LCA affects *S. flexneri* growth under these conditions. At 50 μM, LCA did not alter exponential growth but caused a modest, statistically significant reduction in viability at the stationary phase (**Fig. S2A**) while DCA showed no antimicrobial effect at this concentration (**Fig. S2B**). Thus, LCA triggers aggregation under conditions that impose physiological stress, raising the possibility that aggregation may reflect, in part, a stress-responsive adaptation to LCA. Together, these findings identify LCA as a potent inducer of *S. flexneri* aggregation at physiological concentrations.

### Dissecting the genetic basis of LCA-induced *S. flexneri* aggregation

*S. flexneri* harbors a 218-kb plasmid that encodes essential factors for virulence, including transcriptional regulators VirF, VirB, and MxiE, the type III secretion system (T3SS), and IcsA (39–42) (**Fig. 2A**). To determine whether plasmid-encoded components mediate aggregation in response to LCA, we tested the plasmid-cured strain CFS100 (43). CFS100 failed to aggregate in the presence of 50 or 100 μM LCA (**Fig. 2B** and **S3**), indicating that LCA-induced aggregation requires plasmid-encoded factors.

**Figure 2.**
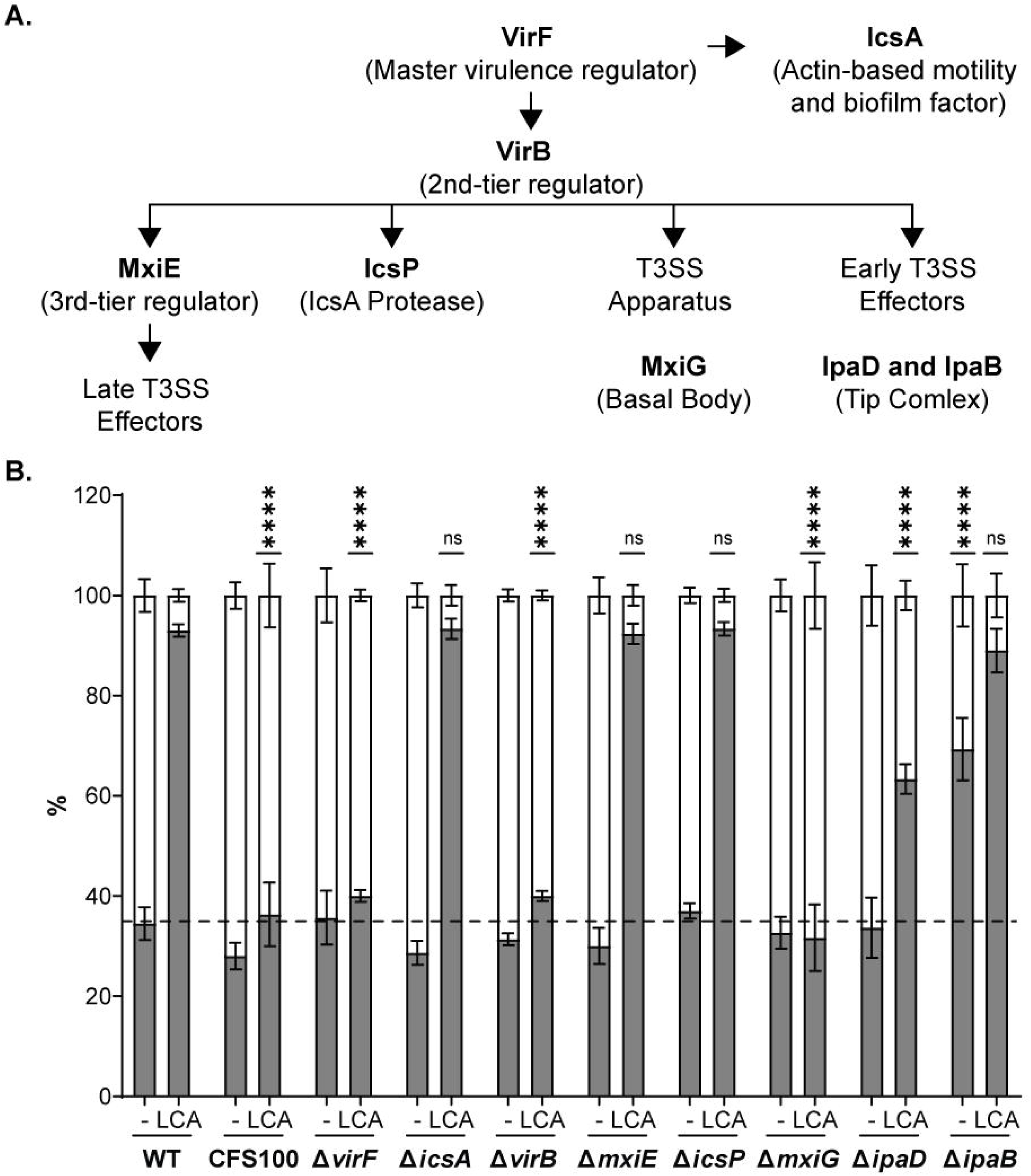
LCA induces S. *f/exneri* aggregation via VirFNirB regulation and T3SS- and lpaD-dependent mechanisms. (A) Schematic of plasmid-encoded regulatory cascades and T3SS-associated genes tested in this study. Bold labels indicate mutated genes; arrows denote established regulatory relationships. (B) Sedimentation assay of S. *flexneri* mutant strains grown in TSB supplemented with 50 **µM** LCA or DMSO vehicle(-). Graph shows mean± SD from three biological repeats; white, planktonic %; gray, aggregate%. Dashed line indicates basal bile acid-independent aggregation. Statistics: one-way ANOVA with Dunnett’s multiple comparison test relative to the corresponding *WT* control. ****, p < 0.0001; ns, non-significant.

We next performed a systematic mutational analysis (**Fig. 2A**) to identify specific genetic determinants. Deletion of *virF*, encoding the master virulence regulator (44), abolished LCA-induced aggregation (Δ*virF*, **Fig. 2B** and **S3**). Among VirF-regulated genes, deletion of *virB* also prevented aggregation, whereas deletion of *icsA* had no effect (ϕ*virB* and ϕ*icsA,* **Fig. 2B** and **S3**). Further analysis of the VirB regulon revealed that deletion of *icsP* or *mxiE* did not impair aggregation (ϕ*icsP* and ϕ*mxiE*, **Fig. 2B** and **S3**). In contrast, deletion of *mxiG*, which encodes a core component of the T3SS basal body (45, 46), completely abolished LCA-induced aggregation (Δ*mxiG*, **Fig. 2B** and **S3**), implicating the T3SS in this process. We then focused on the T3SS tip complex components, IpaD and IpaB effectors (47–49). The Δ*ipaD* mutant exhibited a significant reduction in LCA-induced aggregation (**Fig. 2B** and **S3**), identifying IpaD as a key contributing factor. By contrast, the Δ*ipaB* mutant aggregated normally in the presence of LCA but interestingly showed a hyper-aggregation phenotype in media lacking LCA (**Fig. 2B** and **S3**). This suggests IpaB may negatively regulate basal aggregation when bile acid signals are absent. Consistent with a protein-dependent mechanism, proteinase K treatment eliminated LCA-induced aggregation (**Fig. S4**), supporting the involvement of proteinaceous components, including the T3SS and IpaD. To confirm the specific contribution of IpaD, we constructed an arabinose-inducible *ipaD-HA* expression system and verified its expression and secretion using a Congo red-induced secretion assay (**Fig. 3A** and **3B**). Arabinose induction restored aggregation in the Δ*ipaD* strain (**Fig. 3C**), confirming the functional role of IpaD in LCA-induced aggregation.

**Figure 3.**
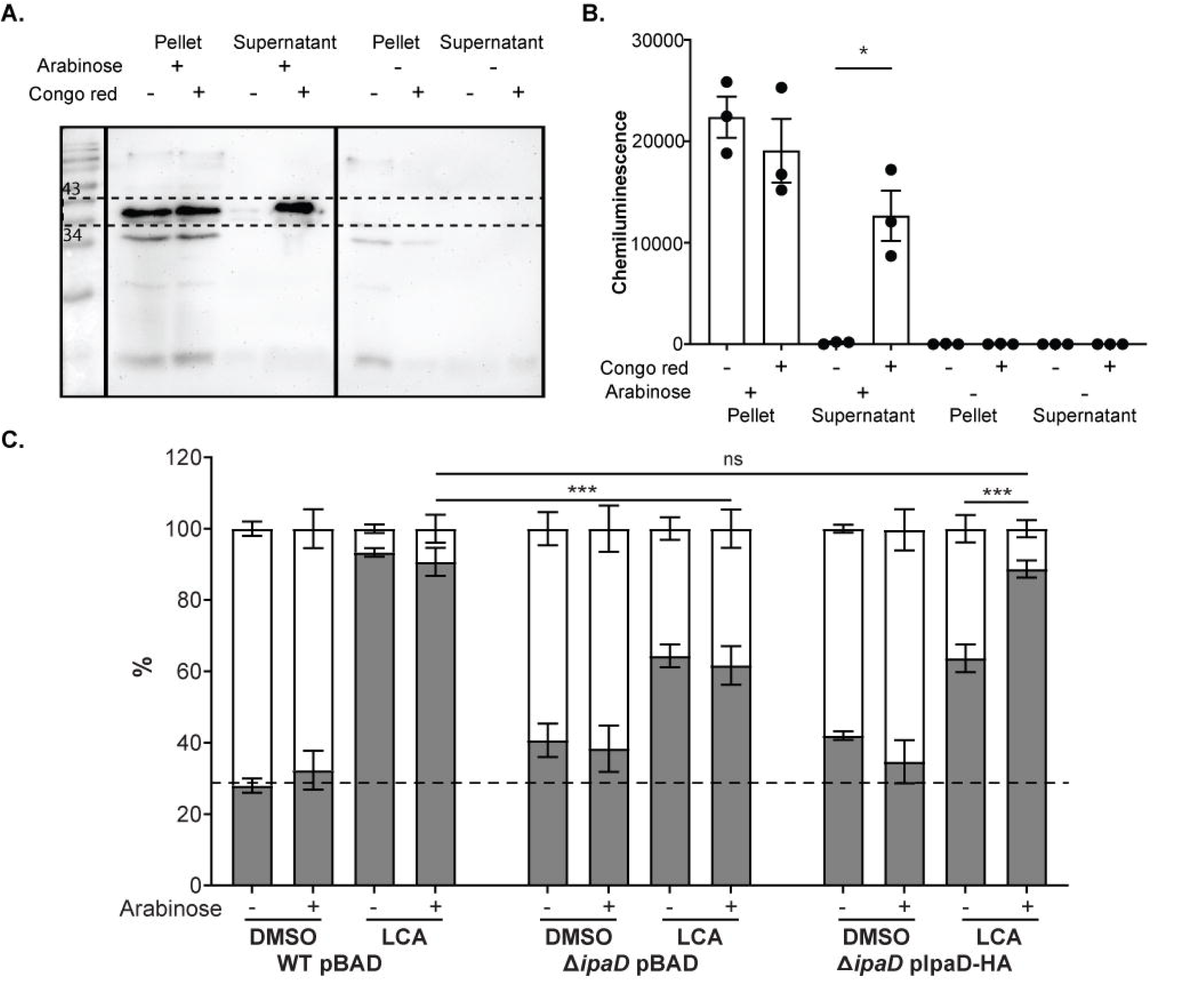
lpaD-HA expression rescues *AipaD* defect in LCA-induced aggregation. (A) Western blot showing arabinose-induced expression and Congo redinduced secretion of lpaD-HA (dashed box: 34-43 kDa) in *fj.ipaD* plpaH-HA. (B) Quantification of lpaD-HA bands. Graph shows mean± SD from three biological repeats. Statistics: paired t-test. *, p < 0.05. (C) Sedimentation assay for LCA-induced aggregation using strains harboring pBAD18 (pBAD) and plpaD-HA. Graph shows mean± SD from three biological repeats; white, planktonic %; gray, aggregate %. Dashed line indicates basal bile acid-independent aggregation. Statistics: one-way ANOVA with Dunnett’s multiple comparison test. ***, p < 0.001; ns, non-significant.

To investigate whether IpaD directly interacts with LCA, we performed *in silico* docking, which identified four potential LCA-contact residues (**Fig. S5A**). However, alanine substitutions at these positions did not alter LCA-induced aggregation relative to IpaD-HA complementation (**Fig. S5B**), suggesting that LCA binding may involve multiple residues or an indirect allosteric contribution of IpaD to aggregation. Together, these findings demonstrate that LCA promotes *S. flexneri* aggregation through a virulence plasmid-encoded, T3SS-dependent mechanism that requires the tip complex protein and effector, IpaD.

### IpaD expression affects the size of LCA-induced *S. flexneri* aggregates

To further elucidate the role of IpaD in LCA-induced aggregation, we analyzed the size of sedimented aggregates formed by the Δ*ipaD* IpaD-HA strain, with or without arabinose-mediated induction of IpaD-HA expression. SYTO9-stained aggregates were imaged and their projected areas quantified. In the absence of arabinose, corresponding to the Δ*ipaD* mutant, sedimented aggregates were significantly smaller, whereas induction of IpaD-HA expression resulted in the formation of larger aggregates (**Fig. 4**). These findings demonstrate that IpaD is required for LCA-induced aggregation and that its expression level influences the size of the resulting sedimented aggregates, supporting its central role in the T3SS-dependent aggregation mechanism.

**Figure 4.**
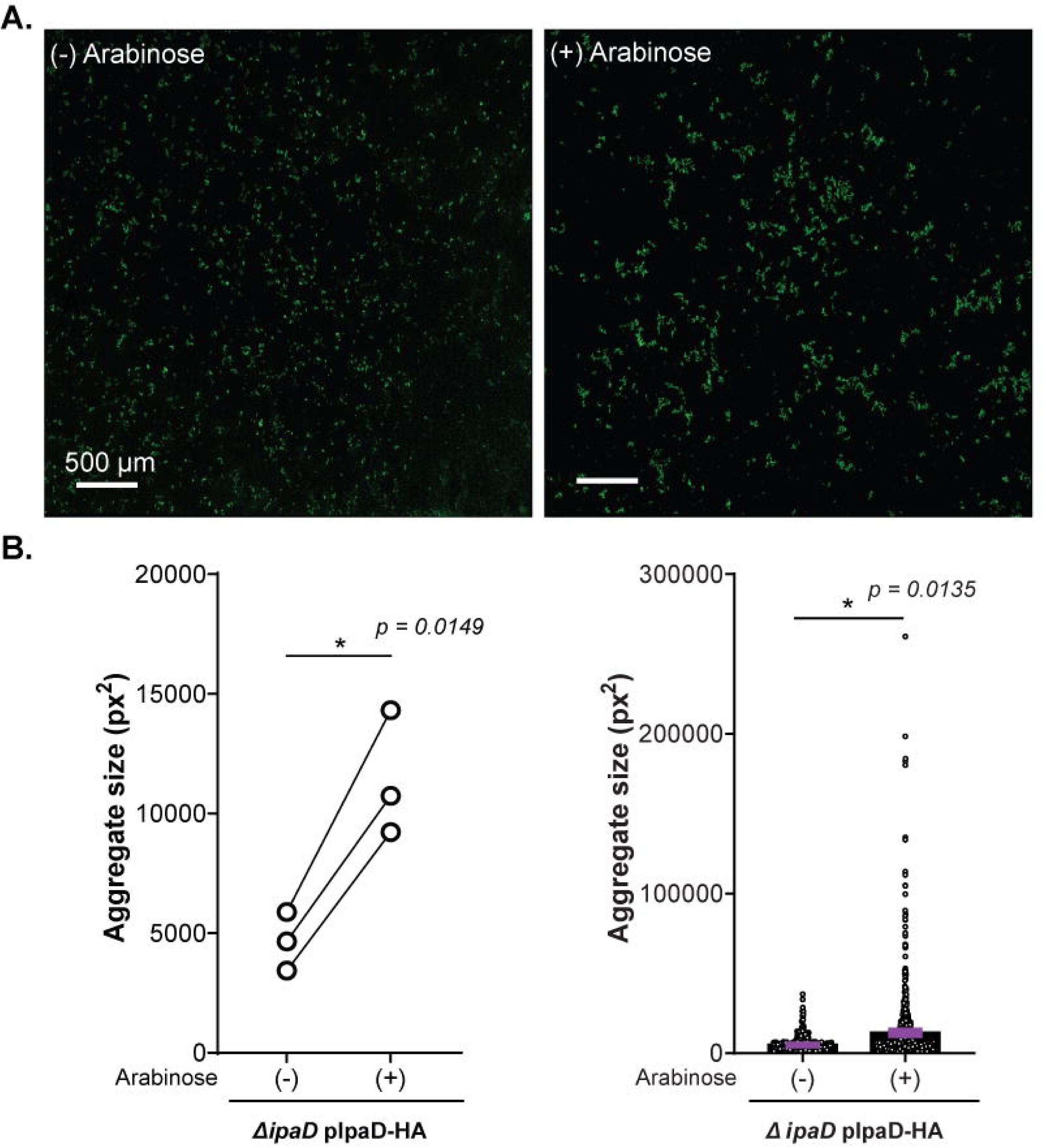
lpaD promotes the formation of large LCA-induced aggregates. (A) Representative widefield images (20x objective) of *t:..ipaD* plpaD-HA sedimented aggregates after growth with 50 **µM** LCA in the absence(-) and presence(+) of 0.1% arabinose, stained with SYTO-9. Scale bar, 500 µm. (B) Quantification of aggregate sizes. Left: Graph shows replicate-level means of projected aggregate area from three biological repeats; each circle represents one repeat, and lines connect paired samples. Statistics: paired t-test. Right: Graph show the distribution of all aggregate areas measured across three biological repeats. Statistics: nested t-test.

### Interactions between LCA-induced *S. flexneri* aggregates and epithelial cells

Given that LCA induces *S. flexneri* aggregation through a T3SS/IpaD-dependent mechanism, we next characterized the interaction between aggregates and colonic epithelial cells during the early phase of infection. To assess adhesion and invasion, we performed gentamicin protection assays using HT-29 cells and quantified adherent and invasive bacteria at 1 hour and 2 hours post-infection, respectively. LCA-induced aggregates produced adherent (**Fig. 5A**) and invasive (**Fig. 5B**) burdens comparable to non-aggregating controls (LCA-independent bacterial pellets), indicating that aggregates retain invasive capacity. To compare early host-pathogen interaction dynamics, we used a T3SS reporter plasmid that constitutively expresses mCherry to label bacteria and produces EGFP upon active T3SS secretion in *S. flexneri* (50). Reporter-expressing non-aggregating bacteria and LCA-induced aggregates were applied to confluent HT-29 monolayers and fixed at 15 and 30 min post-infection, time points selected to capture early aggregate-specific interactions. Infections with LCA-induced aggregates triggered the formation of intense Factin foci by 15 and 30 min, stronger than in infections with non-aggregating controls (**Fig. 6A**). This finding suggests that LCA-induced aggregates manipulate the host cytoskeleton via active secretion of T3SS effectors that modulate actin reorganization dynamics during invasion (51, 52). Consistently, LCA-induced aggregates exhibited higher EGFP/mCherry ratios (EGFP, yellow box, **Fig. 6B**) than non-aggregating controls by 30 min (EGFP, white box, **Fig. 6B**), indicating increased per-bacterium T3SS activity rather than an effect of aggregate size (graph, **Fig. 6B**). Aggregate-associated bacteria were also frequently observed within actin-encased vacuoles (yellow arrows, **Fig. 6B**), an intermediate stage preceding vacuolar escape (53). Together, these data show that LCA-induced aggregates remain invasion-competent and initiate a kinetically different epithelial interaction characterized by robust actin remodeling and early T3SS activation.

**Figure 5.**
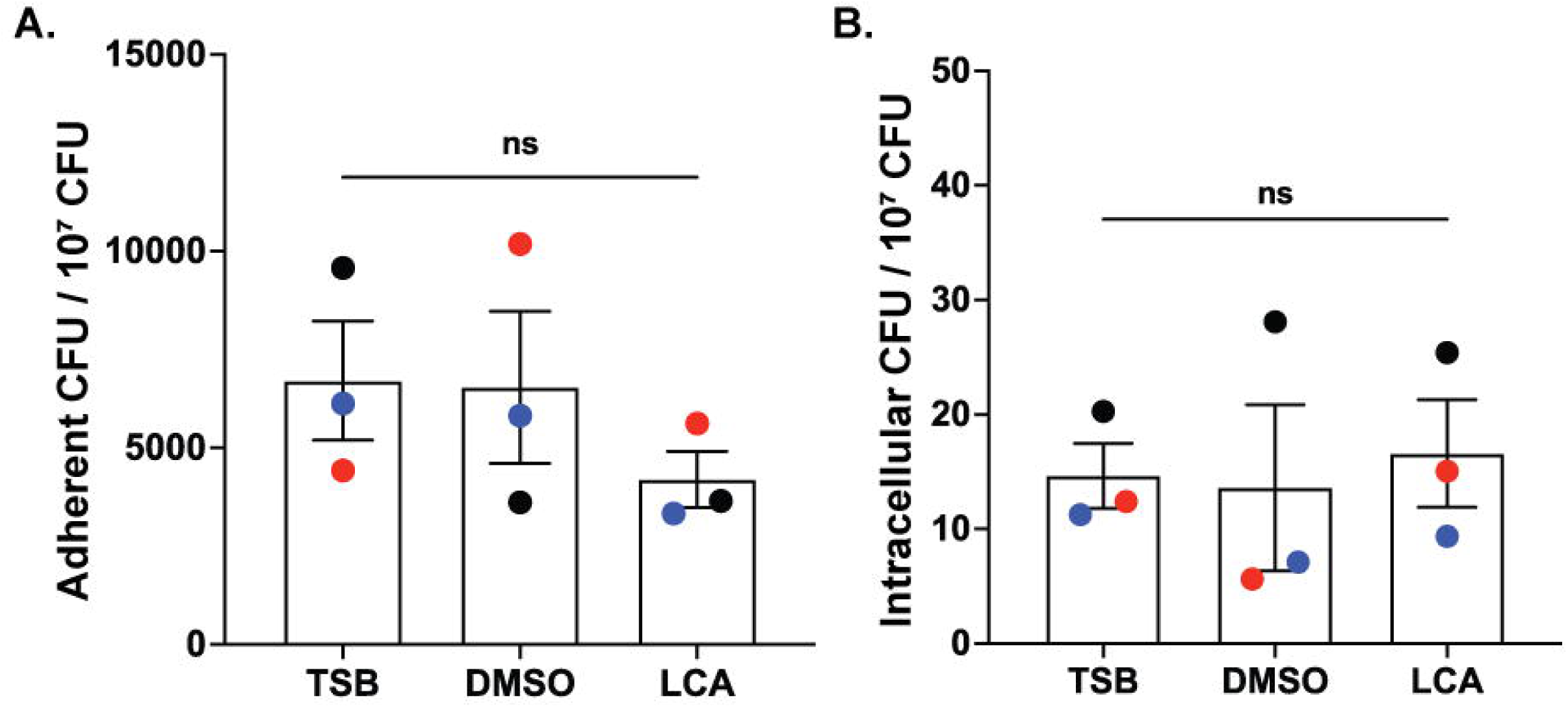
LCA-induced bacterial aggregates adhere to and invade HT-29 cells. (A) Adhesion to HT-29 cells by non-aggregating controls grown in TSB, TSB + vehicle (DMSO), and LCA-induced aggregates (LCA). (B) Invasion of HT-29 cells by non-aggregating controls grown in TSB, TSB + vehicle (DMSO), and LCA-induced aggregates (LCA). Graphs show mean ± SD from three biological repeats; each color represents one biological repeat. Statistics: one-way ANOVA with Tukey’s multiple comparison test. ns, not significant.

**Figure 6.**
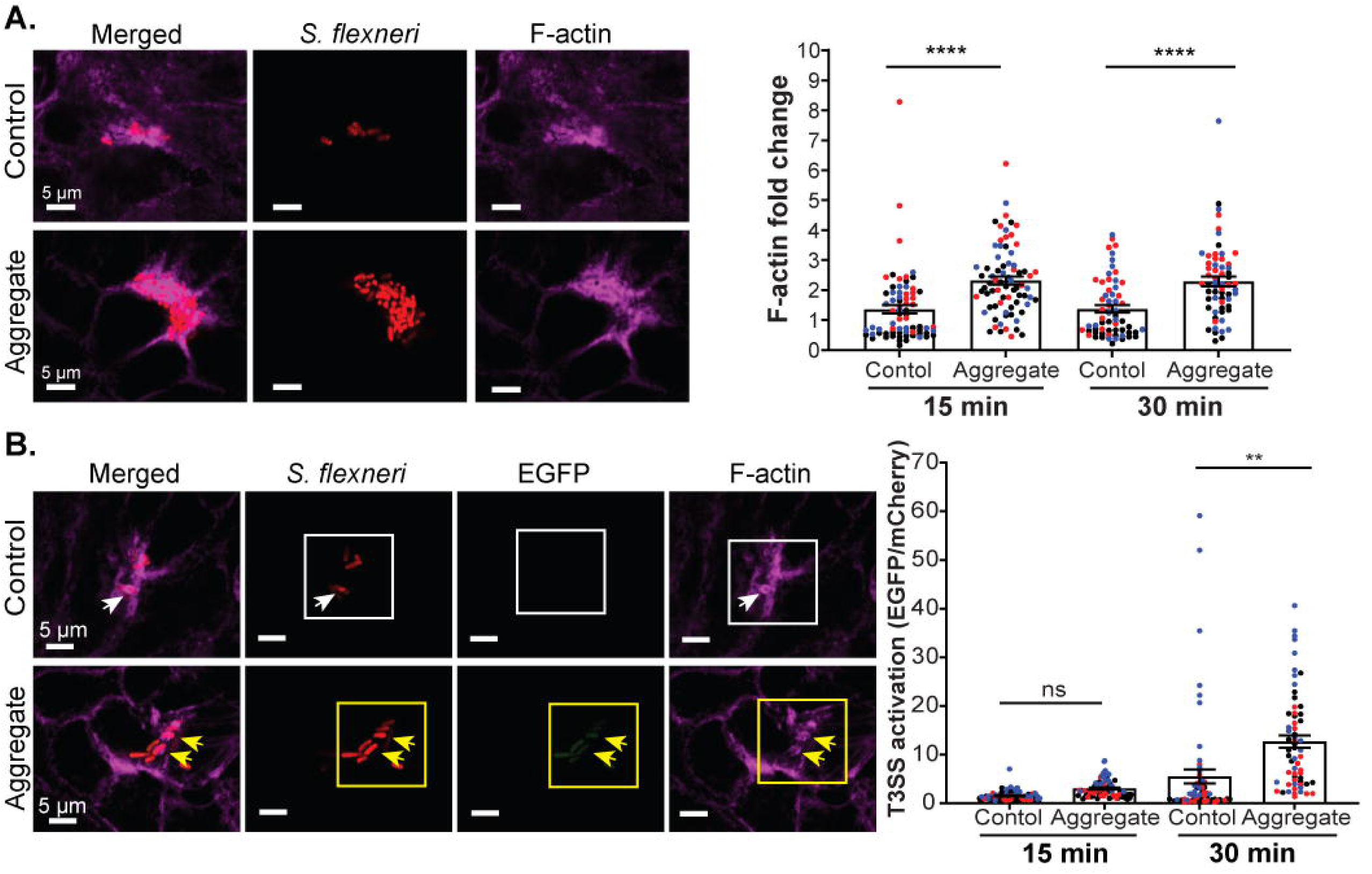
LCA-induced aggregates confer strong actin polymerization and display prompt T3SS activity during cell interactions. (A) Representative images of 15 min infections with S. *f/exneri* grown in TSB containing 0.04% DMSO (Control) or 50 µM LCA (Aggregate) in HT-29 monolayers. Merged image, maximum intensity projection; individual channels, representative single Z-planes; S. *flexneri,* mCherry; F-actin, magenta. Scale bar, 5 µm. Graph shows mean ± SEM and colors indicate three biological repeats. Each point represents individual F-actin focus. Quantification shows F-actin focus intensity normalized to uninfected cell regions. (B) Representative images of 30 min infections with S. *flexneri* grown in TSB containing 0.04% DMSO (Control) or 50 µM LCA (Aggregate) in HT-29 monolayers. Merged image, maximum intensity projection; individual channels, representative single Z-planes; S. *flexneri,* mCherry; Active T3SS, EGFP; F-actin, magenta. White arrows indicate vacuole-associated bacteria; white boxes denote bacteria lacking T3SS activity. Yellow arrows indicate vacuole-associated bacteria; yellow boxes denote T3SS-active bacteria (EGFP-positive). Scale bar, 5 µm. Graph shows mean ± SEM and colors indicate three biological repeats. Each point represents a single measurement. Quantification shows EGFP intensity normalized with associated mCherry intensity to account for bacterial number. Statistics (A) and (B): one-way ANOVA with Dunnett’s test. **, p < 0.01; ****, p < 0.0001; ns, non-significant.

## DISCUSSION

### A specific role for LCA in *S. flexneri* aggregation

We identify the secondary bile acid, LCA as an environmental signal that drives *S. flexneri* aggregation at physiological concentrations (50-100 μM) (54, 55) (**Fig. 1C**), making it far more potent than DCA, which requires substantially higher concentrations to induce aggregation (**Fig. 1D**). LCA does not promote biofilm formation on polystyrene surfaces (**Fig. 1A**), in contrast to DCA, which increases surface-attached biofilm biomass under the same conditions (**Fig. 1B**), as shown previously (33–35). Thus, under these *in vitro* conditions, LCA induces a mode of community assembly distinct from canonical biofilm growth.

LCA exhibits antimicrobial activity (56) and modulates bacterial physiology and virulence factors in other species. For example, it down-regulates toxin expression, toxin activity, and germination in *Clostridioides difficile* and promotes biofilm formation and altered morphology in vancomycin-resistant *Enterococcus faecium* (57–59). Consistent with this broader antimicrobial role, we observed a modest reduction in *S. flexneri* viability in the presence of LCA (**Fig. S2**). These findings support a model in which LCA serves as both a luminal cue and a physiological stressor. It is also possible that LCA-induced aggregation reflects a bacterial response to the antimicrobial stress imposed by this bile acid.

### An extracellular role for IpaD in T3SS-dependent LCA-induced aggregation

Bacterial aggregation can arise through different biophysical mechanisms shaped by environmental constraints on motion or by the presence of bacterial and host-derived polymeric structures (60). Although the molecular mechanism by which LCA promotes aggregation remains unknown, we showed that LCA-induced aggregation requires several virulence plasmid-encoded factors, including the transcriptional regulators VirF and VirB, the T3SS basal body component MxiG, and the T3SS tip protein IpaD (**Fig. 2** and **Fig. S3**), revealing a previously unrecognized, T3SS-dependent mode of community formation triggered by a host-derived metabolite.

Our data identify IpaD as a key contributor to this process. The Δ*ipaD* mutant is impaired in aggregation (**Fig. 2**, **Fig. S3**, and **Fig. 3**), and induction of IpaD expression increases aggregate size (**Fig. 4**), indicating a role for IpaD in the assembly of aggregate architecture. However, IpaD is not the sole determinant of aggregation: the Δ*virF*, Δ*virB*, and Δ*mxiG* mutants are completely defective (**Fig. 2** and **Fig. S3**). This suggests that IpaD functions in concert with other T3SS structural components or early effectors. Late effectors appear dispensable, as the Δ*mxiE* mutant aggregates normally (**Fig. 2** and **Fig. S3**).

IpaD is a T3SS effector essential for invasion of epithelial cells (61). It localizes the tip of the T3SS needle (47), and exposure to DCA allows IpaD to recruit IpaB to the T3SS tip complex (48). In association with IpaB, IpaD also regulates T3SS-dependent secretion by repressing premature effector release (62). We showed that LCA-induced aggregation does not require IpaB, as the Δ*ipaB* strain aggregates normally in the presence of LCA but displays a hyper-aggregation phenotype in the absence of LCA (**Fig. 2B** and **Fig. S3**). These findings suggest that IpaB may function as a negative regulator of bile acid-independent aggregation.

Docking simulations identified four candidate IpaD residues potentially involved in LCA interaction (**Fig. S5A**). Alanine substitutions at these positions do not impair LCA-induced aggregation (**Fig. S5B**). However, given the conservative nature of alanine substitutions, these findings do not rule out the involvement of these residues in LCA-responsive IpaD activity. While direct binding remains to be demonstrated, current evidence suggests that LCA may modulate IpaD function indirectly, possibly by altering its expression, secretion, or interactions with other T3SS components. Taken together, these findings expand the functional repertoire of IpaD beyond its canonical role in invasion, revealing an extracellular, LCA-responsive function in *S. flexneri* aggregation.

### Implications of LCA-induced aggregation for *S. flexneri*-host interactions

Bacterial aggregates are increasingly recognized as pathogenic communities across various infectious diseases (63). These communities can exhibit behaviors distinct from those of planktonic bacteria, particularly during early host-cell interactions. For example, *Streptococcus pyogenes* aggregates, mediated by surface proteins, show enhanced adherence to host cells relative to non-aggregating *S. pyogenes* (64). Similarly, *Pseudomonas aeruginosa* aggregates promote invasion by facilitating their internalization by MDCK cells (65), while *Listeria monocytogenes* aggregates enhance both adhesion and InlB-dependent invasion in HeLa cells (66). These examples illustrate the diverse ways in which aggregates can participate in epithelial invasion. Consistent with invasion-competent aggregates, our gentamicin protection assays show that *S. flexneri* aggregates adhere to and invade epithelial cells at rates comparable to non-aggregating bacteria (**Fig. 5**). However, imaging analyses reveal a distinct phenotype: LCA-induced *S. flexneri* aggregates initiate host cell contact with accelerated early kinetics, characterized by intense actin reorganization and prompt T3SS activation (**Fig. 6**). These findings suggest that, in addition to the well-established invasion mode driven by exponentially growing, non-aggregating *S. flexneri*, LCA-induced aggregate-mediated invasion constitutes an alternative mode for epithelial entry.

When considered with *in vivo* observations of *S. flexneri* aggregates (22, 23), our data support the hypothesis that LCA-induced aggregation serves as a bridge between extracellular colonization and epithelial invasion during *S. flexneri* infection. This mode of entry raises important questions regarding how aggregate-driven invasion affects subsequent intracellular infection dynamics and cellular responses.

Beyond epithelial interactions, LCA-induced aggregates may also confer ecological advantages in the intestinal lumen. First, T3SS/IpaD-dependent aggregation may cluster virulent bacteria in ways that spatially separate them from T3SS-defective subpopulations. Although this possibility remains unexplored, such organization could protect T3SS-proficient cells from luminal factors, such as proteases, that can disrupt T3SS function, thereby helping preserve effector delivery. Second, aggregates may enhance resistance to environmental stressors yet to be characterized. Unlike *S. sonnei, S. flexneri* lacks both a T6SS (67) and capsule (68). We propose that bile acid-driven community formation may help compensate for these deficiencies: DCA primarily promotes biofilm formation (33, 34), whereas LCA drives T3SS/IpaD-dependent aggregation. Together, these distinct bile acid-linked mechanisms may reinforce *S. flexneri* survival in the lumen while simultaneously generating invasion-competent clusters. More broadly, LCA-induced, T3SS-dependent aggregation may reflect a general adaptation among Gram-negative enteric pathogens and a potential target for therapeutic intervention.

## MATERIALS AND METHODS

### Bacterial strains, bile acid media preparation, and cell culture

*Shigella flexneri* strain 2457T, provided by Dr. Benjamin Koestler (Western Michigan University), was used as the model pathogen, and *Escherichia coli* DH5α was used for cloning (**Table 1**). Bacterial stocks were maintained at −80 °C in Tryptic Soy Broth (TSB; Sigma-Aldrich, 22092-500G) supplemented with 20% glycerol. Frozen *S. flexneri* stocks were streaked onto Lysogeny Broth (LB; Fisher, BP1426-2) agar plates containing 10 µg/ml Congo red (Sigma-Aldrich, SDC6277) and incubated at 30°C or 37°C overnight to obtain isolated colonies. When required, culture media were supplemented with ampicillin (100 µg/ml; Sigma-Aldrich Fine Chemicals, NC1004831), kanamycin (30 µg/ml; Thermo Scientific, AC450810100), or chloramphenicol (10 µg/ml; Sigma-Aldrich Fine Chemicals, 501786841).

**Table 1.** Bacterial strains used in this study.

| Strain | Description | Source / Reference |
| --- | --- | --- |
| *S. flexneri* 2457T | Wild type | Dr. Benjamin Koestler, West Michigan University / (41) |
| CFS100 | *pINV*-cured 2457T | Dr. Benjamin Koestler, West Michigan University |
| *S. flexneri virF::kanR* | Insertional mutant | This study |
| *S. flexneri virB::kanR* | Insertional mutant | This study |
| *S. flexneri mxiE::kanR* | Insertional mutant | This study |
| *S. flexneri mxiG::kanR* | Insertional mutant | This study |
| *S. flexneri icsA::kanR* | Insertional mutant | This study |
| *S. flexneri icsP::kanR* | Insertional mutant | This study |
| *S. flexneri ipaD::kanR* | Insertional mutant | This study |
| *S. flexneri* $\Delta ipaD$ | Deletion mutant | This study |
| *S. flexneri ipaB::kanR* | Insertional mutant | This study |
| *E. coli* DH5α | Cloning strain | (34) |

All bile acid solutions were prepared fresh for each experiment. Sodium deoxycholate (DCA; Sigma-Aldrich, D6750-25G) was dissolved directly in TSB and sterilized using 0.22-µm filter units (MilliporeSigma, SLGSR33SS). Lithocholic acid (LCA; Sigma-Aldrich, L6250-10G) was dissolved in dimethyl sulfoxide (DMSO; Fisher Scientific, 317275500ML) and sterilized using DMSO-compatible 0.22-µm filter units (CellTreat, 229757). For growth-curve experiments, bileacid-supplemented TSB was dispensed into 96-well plates after inoculation with bacterial colony suspensions, and optical density at 600 nm (OD_600_) was recorded at 37°C hourly using a microplate reader (Synergy H1, BioTek). Colony-forming units (CFUs) were enumerated at 4 h and 24 h to validate OD_600_ measurements.

The human colorectal epithelial cell line HT-29 (HTB-38; ATCC) was cultured in McCoy’s 5A medium (Gibco, 16600-108) supplemented with 10% heat-inactivated fetal bovine serum (FBS; Gibco, 26140-079) and maintained at 37°C with 5% CO_2_. Upon reaching 70-90% confluency, monolayers were washed three times with phosphate-buffered saline (DPBS; Gibco, 14190-250) and detached with 0.25% trypsin-EDTA. Cells were routinely passaged at 1:5 or 1:10 dilutions and used between passages 4 and 19.

### Construction of *S. flexneri* mutants

Mutant strains were generated using λ-red recombineering (69) (**Table 1**). PCR fragments were amplified from template plasmid pKD4 (**Table 2**), which carries a kanamycin-resistance cassette, using primers listed in **Table 3**. Each fragment contained 42 bp homology regions corresponding to sequences flanking the target gene. *S. flexneri* harboring pKD46 was grown at 30°C with arabinose induction (0.1%) until exponential phase (OD_600_ of 0.5-0.6). Electrocompetent cells were prepared and transformed with PCR products via electroporation (1.8 kV, 400 Ω, 25 µF). After overnight recovery growth, recombinants were selected on kanamycin plates and verified by colony PCR using confirmation primers located outside the deletion region (**Table 3**). To remove the kanamycin cassette, mutants were transformed with pCP20 (**Table 2**), serially passaged at 37 °C, and screened for loss of both kanamycin and ampicillin resistance. Cassette excision was verified by PCR using confirmation primers (**Table 3**). For complementation, IpaD was cloned into pBAD18 (**Table 2**) with a C-terminal HA tag. The gene was amplified by PCR, digested with EcoRI-HF and SphI-HF, and ligated into CIP-treated pBAD18 digested with the same restriction enzymes. Constructs were verified by Sanger sequencing.

**Table 2.** Plasmids and constructs used in this study.

| Plasmid | Description | Source / Reference |
| --- | --- | --- |
| pKD46 | Carries λ-red recombination genes under arabinose induction | NovoPro (V007013) |
| pKD4 | Template plasmid carrying a kanamycin-resistance (*kanR*) cassette flanked by FRT sites | Dr. M. Ammar Zafar, Emory University |
| pCP20 | Carries FLP recombinase for excision of the *kanR* cassette; temperature-sensitive origin | Dr. M. Ammar Zafar, Emory University |
| pTSAR 3.4t | Fluorescent reporter for T3SS activity | Addgene (#210256) |
| pBAD18 | Arabinose inducible gene expression | NovaPro (V001454) |
| pBAD18-IpaD-HA | Complementation plasmid expressing IpaD tagged with HA at the C-terminus | This study |

**Table 3.**
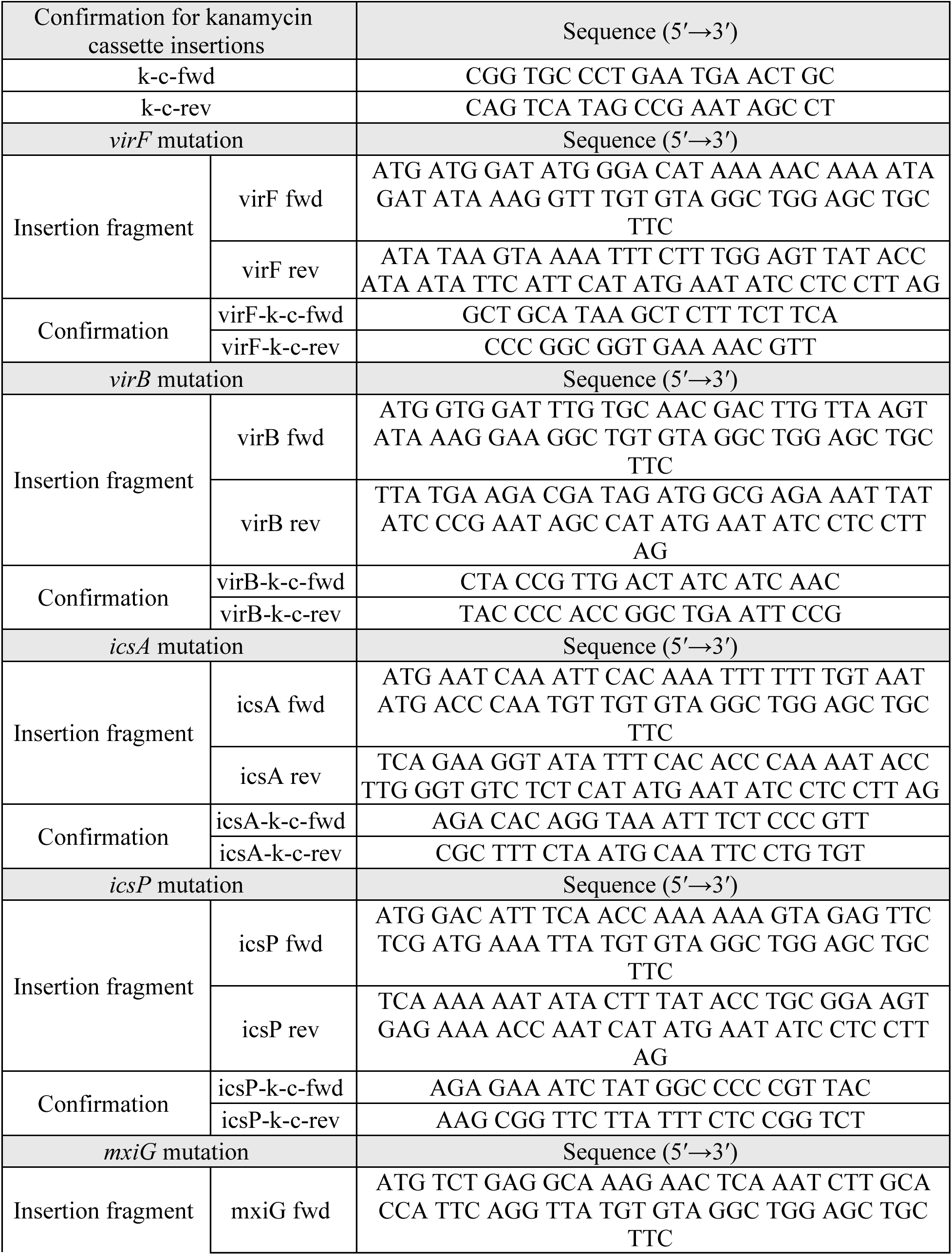
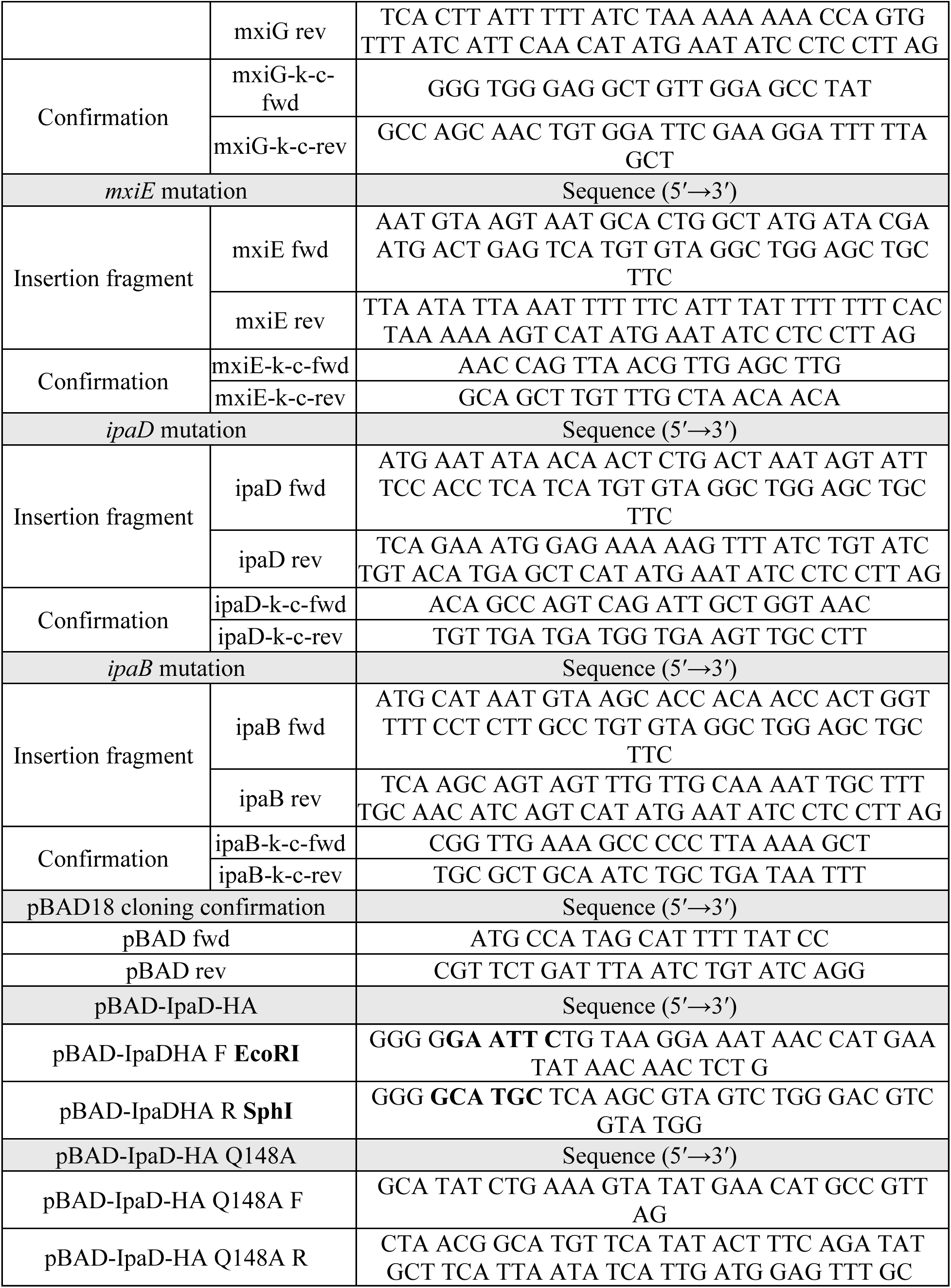
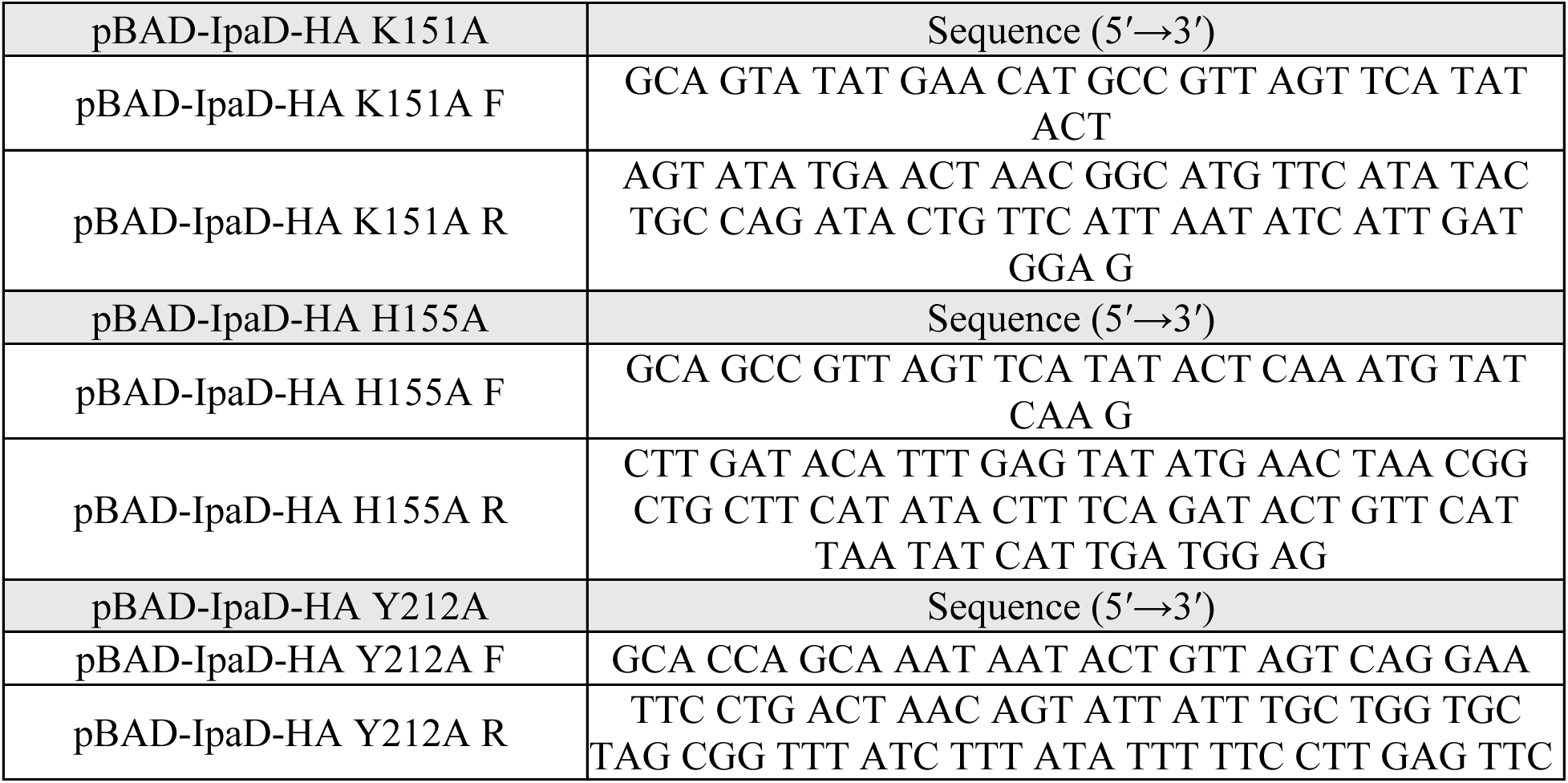
Primers used in this study.

### Sedimentation assay

Isolated *S. flexneri* colonies were resuspended in LB, and 40 µl of suspension was used to inoculate 4 ml TSB supplemented with the indicated LCA or DCA concentrations in 15 ml culture tubes. Cultures were incubated statically at 37°C for 24 h. TSB alone or TSB supplemented with DMSO served as controls that exhibit bile acid-independent basal aggregation or bacterial pellet, hereafter referred to as the non-aggregating control. After incubation, a 150 µl aliquot was collected from the culture surface (planktonic-enriched fraction). The remaining culture was then vortexed to fully resuspend aggregates, and a separate 150 µl aliquot was taken as the total culture. OD_600_ values were measured for both the planktonic and total fractions using the microplate reader (Synergy H1, BioTek), and relative aggregation percentage was calculated as:

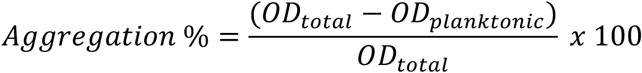

Samples for CFU determination from planktonic and aggregated populations were obtained from cultures grown in parallel under the same conditions used for OD_600_ measurements. From each culture, 100 µL of the planktonic fraction was collected from the surface and plated to LB agar supplemented with Congo red. Additionally, 100 µl of the aggregate fraction was fully resuspended and plated. For the non-aggregating control, 100 µl of the bacterial pellet -representing bile acid-independent background sediment- was collected and plated using the same procedure. CFU-based aggregation ratios were calculated using the same approach as OD_600_-based aggregation ratios, following the formula provided. For proteinase K treatment, culture media were supplemented with 100 µg/mL proteinase K (QIAGEN, 19131) prior to bacterial inoculation.

### Secretion assay

IpaD-HA secretion by the T3SS was induced with Congo red as described previously (70). Briefly, 2 ml of exponential phase cultures were pelleted (761 g, 10 min) and resuspended in 500 µl sterile PBS with and without 0.05% (wt/vol) Congo red. Following incubation at 37°C for 1 h, samples were centrifuged (16100 g, 30 s) to separate supernatants from pellets. Supernatants were precipitated with 10% trichloroacetic acid (Sigma-Aldrich, T0699-100ML) on ice for 1 h. Bacterial pellets were resuspended in 500 µl PBS and mixed 1:1 with 2X Laemmli sample buffer (Bio-Rad, 1610737) containing 0.1 M dithiothreitol (Fisher Scientific, FERR0861), followed by boiling for 10 min. Proteins were resolved on 15% SDS-polyacrylamide gels, transferred to PVDF membranes (BioRad, 1620177), and blocked in 5% bovine serum albumin (BSA; Fisher BioReagents, BP1600-100) in PBS containing 0.1% Tween 20 (PBST) (Sigma-Aldrich, P7949-100ML) for 1 h at room temperature. Membranes were incubated overnight at 4°C with anti-HA monoclonal antibody (Proteintech, 66006-2-Ig; 1:10000) in 5% BSA-PBST, washed, and probed with HRP-conjugated goat anti-mouse IgG (BioRad, 1706516; 1:50000) for 1 h at room temperature. Protein bands were visualized using Pierce™ ECL Western Blotting Substrate (Fisher Scientific, PI32209) and imaged with a Bio-Rad ChemiDoc system. ImageJ was used for quantification.

### Confocal imaging of aggregates

Preparation and inoculation for LCA-supplemented cultures followed the 24 h sedimentation assay, with an additional 0.1% arabinose condition to induce expression of plasmid-borne IpaD-HA. A 50 µl aliquot was withdrawn from the aggregate layer using a wide-bore pipette tip and transferred into 500 µl of PBS containing SYTO9 (ThermoFisher Scientific, S34854; 1:1000). Samples were incubated for 15 min at room temperature, transferred into 8-well µ-Slides (Ibidi, 80806), and allowed to settle for 20 min at 37 °C. The staining solution was then aspirated and replaced with fresh PBS containing SYTO9 to maintain fluorescence. Images were acquired using a Leica STELLARIS DMI8 confocal microscope equipped with a 20X water-immersion objective. Image analysis was performed using MetaXpress 6 (Molecular Devices). Thresholds were uniformly applied across all images to remove background signal and noise. Aggregate areas were quantified using the Integrated Morphometry Analysis module.

### Aggregate adherence and invasion in HT-29 cell monolayers

After 24 h of static growth in TSB supplemented with 50 µM LCA or 0.04% DMSO (control), 50 µl of the aggregate fraction or the control bacterial pellet -representing LCA-independent basal aggregation used as the non-aggregating control- was collected using a manually cut pipette tip. Samples were applied to 4-day confluent HT-29 monolayers in 24-well plates (Fisher Scientific, NC1713666) containing 500 µl of fresh McCoy’s 5A medium supplemented with 10% heat-inactivated FBS. Plates were centrifuged for 2 min at 800 rpm to promote contact between bacteria and epithelial cells, then incubated at 37°C with 5% CO_2_. For adherence assays, monolayers were incubated for 1 h before washing three times with PBS to remove non-adherent bacteria. For invasion assays, parallel wells were incubated for 1 h to allow adherence, then the medium was removed and replaced with medium containing 50 µg/ml gentamicin (Sigma-Aldrich, NC0363642). Plates were incubated in CO_2_ incubator at 37°C for 1 h. After incubation, monolayers were washed three times with DPBS and lysed with ice-cold 0.1% Triton X-100 (Bio-Rad, 1610407) for 12 min to release intracellular bacteria. Lysates were serially diluted and plated on LB agar plates supplemented with Cong red to enumerate CFUs. Percent adhesion was calculated as the number of CFUs recovered after the 1-hour infection (adherence step) divided by the total CFUs in the corresponding inoculum (aggregate or non-aggregating control). Percent invasion was calculated as the number of CFUs recovered after the full 2-hour infection period divided by the total CFUs in the corresponding inoculum.

### Confocal imaging of aggregate-cell interactions

*S. flexneri* strains harboring the pTSAR 3.4t plasmid (50), which constitutively express mCherry and produce EGFP upon T3SS activation, were grown for 24 h under static conditions in TSB supplemented with 50 µM LCA or 0.04% DMSO (control). The LCA-induced aggregate fraction, or the bacterial pellet from the non-aggregating control, was collected with a wide-bore pipette tip and applied to confluent HT-29 monolayers grown on 1.5 glass coverslips (Fisher Scientific, NC1418755) at an MOI of 1. Plates were centrifuged for 2 min at 800 rpm to facilitate contact between bacteria and epithelial cells, then incubated at 37 °C with 5% CO_2_ for 15 or 30 min. After incubation, coverslips were removed, immediately fixed with 4% paraformaldehyde (Fisher Scientific, 50-980-487) for 15 min, and washed three times with DPBS. Fixed samples were incubated with Alexa Fluor-680 Phalloidin (ThermoFisher Scientific, A22286; 1:1000) to visualize F-actin and assess ruffling at bacteria-cell contact sites. Coverslips were mounted with ProLong Glass Antifade Mountant (ThermoFisher Scientific, P36980), and images were acquired using a Leica STELLARIS DMI8 confocal microscope equipped with a 63X oil-immersion objective. In each experiment, 20-25 infection foci were randomly selected across each coverslip to avoid sampling bias. Image analysis was performed using LAS X software (Leica) to quantify: (i) the area of individual F-actin foci at aggregate-cell or non-aggregating control-cell interfaces, (ii) the total mCherry fluorescence associated with each focus (representing either LCA-induced aggregates or the basal, LCA-independent bacterial pellet used as the non-aggregating control), and (iii) the corresponding total EGFP fluorescence intensity to assess T3SS activation. Normalized T3SS activity was calculated as EGFP/mCherry for each focus, thereby correcting for bacterial number and enabling direct comparison of T3SS activity across aggregate and non-aggregating control conditions.

### In silico docking

The IpaD protein from *S. flexneri* was modeled using AlphaFold2 implemented via the ColabFold platform (71, 72). ColabFold was run on the DEAC high-performance computing cluster using default parameters, with no custom templates or distance restraints provided. Five models were generated, and the highest-confidence prediction (ranked by predicted LDDT) was selected as the receptor structure for docking. The selected AlphaFold2 model was prepared and converted to PDBQT format using AutoDock Tools (ADT) v.1.5.4 (73). The 3D structure of LCA (PubChem Compound CID: 9903) was retrieved from the PubChem database in SDF format (74). Ligand preparation was performed with ADT; all hydrogen atoms were added, and Gasteiger partial charges were assigned to the LCA molecule, and non-polar hydrogens were merged into a PDBQT file. Molecular docking of LCA to IpaD was carried out using AutoDock Vina (v. 1.5.2) (75). The docking search space was defined as a grid box of 50 Å per side, centered on the known deoxycholic acid (DCA) binding site of IpaD. All docking runs used the default Vina scoring function and search parameters and were performed on the DEAC cluster. The top nine binding poses were generated by Vina, and the highest-ranked pose (lowest predicted binding free energy) was selected for further analysis. AutoDock Vina was run in default exhaustiveness mode, and results were analyzed using VMD(v.1.9.4) (76).

### Statistical analysis

All experiments were performed with at least three independent biological replicates. Statistical analyses were conducted using GraphPad Prism 10 (GraphPad Software). Nested t-tests were used for aggregate area quantifications to account for variability among independent experiments. Unpaired two-tailed Student’s t-tests were used for pairwise comparisons of aggregate size measurements. One-way ANOVA with Dunnett’s comparison test was applied for sedimentation assays, F-actin foci size measurements, and T3SS activity quantifications and Tukey’s test for adherence/invasion assays.

## ACKNOWLEDGEMENTS

We thank Dr. Benjamin Koestler for providing *Shigella flexneri* 2457T and CFS100 strains and Dr. M. Ammar Zafar for the generous gift of λ-red recombineering plasmids. We are grateful to Dr. Jörn Coers and members of his laboratory for their valuable feedback during manuscript preparation, and to Dr. Martha Alexander-Miller for her thoughtful input on the manuscript. Portions of the computations were performed on the Wake Forest University DEAC Cluster, a centrally managed resource with support provided in part by Wake Forest University. We also acknowledge the Wake Forest Baptist Comprehensive Cancer Center Crystallography & Computational Biosciences Shared Resource, supported by the National Cancer Institute’s Cancer Center Support Grant (award number P30CA012197). The content is solely the responsibility of the authors and does not necessarily represent the official views of the National Cancer Institute. This work was supported by start-up funding from Wake Forest University School of Medicine to Dr. Volkan K. Köseoğlu.

## REFERENCES

1. Kotloff KL, Riddle MS, Platts-Mills JA, Pavlinac P, Zaidi AKM. 2018. Shigellosis. The Lancet 391:801–812.

2. Christie AB. 1968. Bacillary dysentery. Br Med J 2:285–288.

3. Khalil IA, Troeger C, Blacker BF, Rao PC, Brown A, Atherly DE, Brewer TG, Engmann CM, Houpt ER, Kang G, Kotloff KL, Levine MM, Luby SP, MacLennan CA, Pan WK, Pavlinac PB, Platts-Mills JA, Qadri F, Riddle MS, Ryan ET, Shoultz DA, Steele AD, Walson JL, Sanders JW, Mokdad AH, Murray CJL, Hay SI, Reiner RC. 2018. Morbidity and mortality due to shigella and enterotoxigenic Escherichia coli diarrhoea: the Global Burden of Disease Study 1990–2016. The Lancet Infectious Diseases 18:1229–1240.

4. Rogawski ET, Liu J, Platts-Mills JA, Kabir F, Lertsethtakarn P, Siguas M, Khan SS, Praharaj I, Murei A, Nshama R, Mujaga B, Havt A, Maciel IA, Operario DJ, Taniuchi M, Gratz J, Stroup SE, Roberts JH, Kalam A, Aziz F, Qureshi S, Islam MO, Sakpaisal P, Silapong S, Yori PP, Rajendiran R, Benny B, McGrath M, Seidman JC, Lang D, Gottlieb M, Guerrant RL, Lima AAM, Leite JP, Samie A, Bessong PO, Page N, Bodhidatta L, Mason C, Shrestha S, Kiwelu I, Mduma ER, Iqbal NT, Bhutta ZA, Ahmed T, Haque R, Kang G, Kosek MN, Houpt ER, Acosta AM, Rios de Burga R, Chavez CB, Flores JT, Olotegui MP, Pinedo SR, Trigoso DR, Vasquez AO, Ahmed I, Alam D, Ali A, Rasheed M, Soofi S, Turab A, Yousafzai A, Zaidi AK, Shrestha B, Rayamajhi BB, Strand T, Ammu G, Babji S, Bose A, George AT, Hariraju D, Jennifer MS, John S, Kaki S, Karunakaran P, Koshy B, Lazarus RP, Muliyil J, Ragasudha P, Raghava MV, Raju S, Ramachandran A, Ramadas R, Ramanujam K, Rose A, Roshan R, Sharma SL, Sundaram S, Thomas RJ, Pan WK, Ambikapathi R, Carreon JD, Doan V, Hoest C, Knobler S, Miller MA, Psaki S, Rasmussen Z, Richard SA, Tountas KH, Svensen E, Amour C, Bayyo E, Mvungi R, Pascal J, Yarrot L, Barrett L, Dillingham R, Petri WA, Scharf R, Ahmed AS, Alam MA, Haque U, Hossain MI, Islam M, Mahfuz M, Mondal D, Nahar B, Tofail F, Chandyo RK, Shrestha PS, Shrestha R, Ulak M, Bauck A, Black R, Caulfield L, Checkley W, Lee G, Schulze K, Scott S, Murray-Kolb LE, Ross AC, Schaefer B, Simons S, Pendergast L, Abreu CB, Costa H, Di Moura A, Filho JQ, Leite ÁM, Lima NL, Lima IF, Maciel BL, Medeiros PH, Moraes M, Mota FS, Oriá RB, Quetz J, Soares AM, Mota RM, Patil CL, Mahopo C, Maphula A, Nyathi E. 2018. Use of quantitative molecular diagnostic methods to investigate the effect of enteropathogen infections on linear growth in children in low-resource settings: longitudinal analysis of results from the MAL-ED cohort study. The Lancet Global Health 6:e1319–e1328.

5. Nasrin D, Blackwelder WC, Sommerfelt H, Wu Y, Farag TH, Panchalingam S, Biswas K, Saha D, Jahangir Hossain M, Sow SO, Reiman RFB, Sur D, Faruque ASG, Zaidi AKM, Sanogo D, Tamboura B, Onwuchekwa U, Manna B, Ramamurthy T, Kanungo S, Omore R, Ochieng JB, Oundo JO, Das SK, Ahmed S, Qureshi S, Quadri F, Adegbola RA, Antonio M, Mandomando I, Nhampossa T, Bassat Q, Roose A, O’Reilly CE, Mintz ED, Ramakrishnan U, Powell H, Liang Y, Nataro JP, Levine MM, Kotloff KL. 2021. Pathogens Associated With Linear Growth Faltering in Children With Diarrhea and Impact of Antibiotic Treatment: The Global Enteric Multicenter Study. The Journal of Infectious Diseases 224:S848–S855.

6. Livio S, Strockbine NA, Panchalingam S, Tennant SM, Barry EM, Marohn ME, Antonio M, Hossain A, Mandomando I, Ochieng JB, Oundo JO, Qureshi S, Ramamurthy T, Tamboura B, Adegbola RA, Hossain MJ, Saha D, Sen S, Faruque ASG, Alonso PL, Breiman RF, Zaidi AKM, Sur D, Sow SO, Berkeley LY, O’Reilly CE, Mintz ED, Biswas K, Cohen D, Farag TH, Nasrin D, Wu Y, Blackwelder WC, Kotloff KL, Nataro JP, Levine MM. 2014. Shigella Isolates From the Global Enteric Multicenter Study Inform Vaccine Development. Clinical Infectious Diseases 59:933–941.

7. Musher DM, Musher BL. 2004. Contagious Acute Gastrointestinal Infections. The New England Journal of Medicine 351:2466.

8. Mani S, Wierzba T, Walker RI. 2016. Status of vaccine research and development for *Shigella*. Vaccine 34:2887–2894.

9. Frost I, Sati H, Garcia-Vello P, Hasso-Agopsowicz M, Lienhardt C, Gigante V, Beyer P. 2023. The role of bacterial vaccines in the fight against antimicrobial resistance: an analysis of the preclinical and clinical development pipeline. The Lancet Microbe 4:e113–e125.

10. WHO bacterial priority pathogens list, 2024: Bacterial pathogens of public health importance to guide research, development and strategies to prevent and control antimicrobial resistance. https://www.who.int/publications-detail-redirect/9789240093461. Retrieved 20 May 2024.

11. Kotloff KL, Winickoff JP, Ivanoff B, Clemens JD, Swerdlow DL, Sansonetti PJ, Adak GK, Levine MM. 1999. Global burden of Shigella infections: implications for vaccine development and implementation of control strategies. Bull World Health Organ 77:651–666.

12. Labrec EH, Schneider H, Magnani TJ, Formal SB. 1964. Epithelial cell penetration as an essential step in the pathogenesis of bacillary dysentery. Journal of Bacteriology 88:1503–1518.

13. Takeuchi A, Formal SB, Sprinz H. 1968. Exerimental acute colitis in the Rhesus monkey following peroral infection with Shigella flexneri. An electron microscope study. Am J Pathol 52:503–529.

14. Anand BS, Malhotra V, Bhattacharya SK, Datta P, Datta D, Sen D, Bhattacharya MK, Mukherjee PP, Pal SC. 1986. Rectal Histology in Acute Bacillary Dysentery. Gastroenterology 90:654–660.

15. Mathan MM, Mathan VI. 1991. Morphology of Rectal Mucosa of Patients with Shigellosis. Reviews of Infectious Diseases 13:S314–S318.

16. Carayol N, Nhieu GTV. 2013. The Inside Story of Shigella Invasion of Intestinal Epithelial Cells. Cold Spring Harb Perspect Med 3:a016717.

17. Bernardini ML, Mounier J, d’Hauteville H, Coquis Rondon M, Sansonetti PJ. 1989. Identification of icsA, a plasmid locus of Shigella flexneri that governs bacterial intra- and intercellular spread through interaction with F-actin. Proceedings of the National Academy of Sciences 86:3867–3871.

18. Goldberg MB, Bârzu O, Parsot C, Sansonetti PJ. 1993. Unipolar localization and ATPase activity of IcsA, a Shigella flexneri protein involved in intracellular movement. Journal of Bacteriology 175:2189–2196.

19. Agaisse H. 2016. Molecular and Cellular Mechanisms of Shigella flexneri Dissemination. Frontiers in Cellular and Infection Microbiology 6.

20. Raab JE, Hamilton DJ, Harju TB, Huynh TN, Russo BC. 2024. Pushing boundaries: mechanisms enabling bacterial pathogens to spread between cells. Infection and Immunity 0:e00524–23.

21. Chai SJ, Gu W, O’Connor KA, Richardson LC, Tauxe RV. 2019. Incubation periods of enteric illnesses in foodborne outbreaks, United States, 1998–2013. Epidemiology & Infection 147:e285.

22. Kuehl CJ, D’Gama JD, Warr AR, Waldor MK. 2020. An Oral Inoculation Infant Rabbit Model for Shigella Infection. mBio 11:e03105–19.

23. Q.S. Medeiros PH, Ledwaba SE, Bolick DT, Giallourou N, Yum LK, Costa DVS, Oriá RB, Barry EM, Swann JR, Lima AÂM, Agaisse H, Guerrant RL. 2019. A murine model of diarrhea, growth impairment and metabolic disturbances with Shigella flexneri infection and the role of zinc deficiency. Gut Microbes 10:615–630.

24. Sistrunk JR, Nickerson KP, Chanin RB, Rasko DA, Faherty CS. 2016. Survival of the Fittest: How Bacterial Pathogens Utilize Bile To Enhance Infection. Clinical Microbiology Reviews 29:819–836.

25. Hofmann AF. 1999. Bile Acids: The Good, the Bad, and the Ugly. Physiology 14:24–29.

26. Macierzanka A, Torcello-Gómez A, Jungnickel C, Maldonado-Valderrama J. 2019. Bile salts in digestion and transport of lipids. Advances in Colloid and Interface Science 274:102045.

27. Dawson PA, Karpen SJ. 2015. Intestinal transport and metabolism of bile acids. Journal of Lipid Research 56:1085–1099.

28. Ridlon JM, Kang D-J, Hylemon PB. 2006. Bile salt biotransformations by human intestinal bacteria. Journal of Lipid Research 47:241–259.

29. B Zhu C, Fuchs CD, Halilbasic E, Trauner M. 2016 Bile acids in regulation of inflammation and immunity: friend or foe? Clin Exp Rheumatol 34(4 Suppl 98):25-31.025.

30. Ciaula AD, Garruti G, Baccetto RL, Molina-Molina E, Bonfrate L, Portincasa P, Wang DQH. 2018. Bile Acid Physiology. Ann Hepatol 16:4–14.

31. Zhou H, Hylemon PB. 2014. Bile acids are nutrient signaling hormones. Steroids 86:62–68.

32. Larabi AB, Masson HLP, Bäumler AJ. 2023. Bile acids as modulators of gut microbiota composition and function. Gut Microbes 15:2172671.

33. Nickerson KP, Chanin RB, Sistrunk JR, Rasko DA, Fink PJ, Barry EM, Nataro JP, Faherty CS. 2017. Analysis of Shigella flexneri Resistance, Biofilm Formation, and Transcriptional Profile in Response to Bile Salts. Infection and Immunity 85:10.1128/iai.01067-16.

34. Köseoğlu VK, Hall CP, Rodríguez-López EM, Agaisse H. 2019. The Autotransporter IcsA Promotes Shigella flexneri Biofilm Formation in the Presence of Bile Salts. Infection and Immunity 87:10.1128/iai.00861-18.

35. Chiang I-L, Wang Y, Fujii S, Muegge BD, Lu Q, Tarr PI, Stappenbeck TS. 2021. Biofilm Formation and Virulence of Shigella flexneri Are Modulated by pH of Gastrointestinal Tract. Infection and Immunity 89.

36. Pope LM, Reed KE, Payne SM. 1995. Increased protein secretion and adherence to HeLa cells by Shigella spp. following growth in the presence of bile salts. Infection and Immunity 63:3642–3648.

37. Faherty CS, Redman JC, Rasko DA, Barry EM, Nataro JP. 2012. Shigella flexneri effectors OspE1 and OspE2 mediate induced adherence to the colonic epithelium following bile salts exposure. Mol Microbiol 85:107–121.

38. Brotcke Zumsteg A, Goosmann C, Brinkmann V, Morona R, Zychlinsky A. 2014. IcsA Is a Shigella flexneri Adhesin Regulated by the Type III Secretion System and Required for Pathogenesis. Cell Host & Microbe 15:435–445.

39. Buchrieser C, Glaser P, Rusniok C, Nedjari H, D’Hauteville H, Kunst F, Sansonetti P, Parsot C. 2000. The virulence plasmid pWR100 and the repertoire of proteins secreted by the type III secretion apparatus of Shigella flexneri. Mol Microbiol 38:760–771.

40. Venkatesan MM, Goldberg MB, Rose DJ, Grotbeck EJ, Burland V, Blattner FR. 2001. Complete DNA Sequence and Analysis of the Large Virulence Plasmid of Shigella flexneri. Infection and Immunity 69:3271–3285.

41. Wei J, Goldberg MB, Burland V, Venkatesan MM, Deng W, Fournier G, Mayhew GF, Plunkett G, Rose DJ, Darling A, Mau B, Perna NT, Payne SM, Runyen-Janecky LJ, Zhou S, Schwartz DC, Blattner FR. 2003. Complete genome sequence and comparative genomics of Shigella flexneri serotype 2a strain 2457T. Infect Immun 71:2775–2786.

42. Dorman CJ. 2004. Virulence Gene Regulation in Shigella. EcoSal Plus 1:10.1128/ecosalplus.8.9.3.

43. Marman HE, Mey AR, Payne SM. 2014. Elongation Factor P and Modifying Enzyme PoxA Are Necessary for Virulence of Shigella flexneri. Infection and Immunity 82:3612–3621.

44. Di Martino ML, Falconi M, Micheli G, Colonna B, Prosseda G. 2016. The Multifaceted Activity of the VirF Regulatory Protein in the Shigella Lifestyle. Frontiers in Molecular Biosciences 3.

45. Allaoui A, Sansonetti PJ, Ménard R, Barzu S, Mounier J, Phalipon A, Parsot C. 1995. MxiG, a membrane protein required for secretion of Shigella spp. Ipa invasins: involvement in entry into epithelial cells and in intercellular dissemination. Molecular Microbiology 17:461–470.

46. Blocker A, Jouihri N, Larquet E, Gounon P, Ebel F, Parsot C, Sansonetti P, Allaoui A. 2001. Structure and composition of the Shigella flexneri‘needle complex’, a part of its type III secreton. Molecular Microbiology 39:652–663.

47. Espina M, Olive AJ, Kenjale R, Moore DS, Ausar SF, Kaminski RW, Oaks EV, Middaugh CR, Picking WD, Picking WL. 2006. IpaD Localizes to the Tip of the Type III Secretion System Needle of Shigella flexneri. Infection and Immunity 74:4391–4400.

48. Olive AJ, Kenjale R, Espina M, Moore DS, Picking WL, Picking WD. 2007. Bile Salts Stimulate Recruitment of IpaB to the Shigella flexneri Surface, Where It Colocalizes with IpaD at the Tip of the Type III Secretion Needle. Infection and Immunity 75:2626–2629.

49. Picking WL, Picking WD. 2016. The Many Faces of IpaB. Front Cell Infect Microbiol 6.

50. Campbell-Valois F-X, Schnupf P, Nigro G, Sachse M, Sansonetti PJ, Parsot C. 2014. A Fluorescent Reporter Reveals On/Off Regulation of the Shigella Type III Secretion Apparatus during Entry and Cell-to-Cell Spread. Cell Host & Microbe 15:177–189.

51. Carayol N, Tran Van Nhieu G. 2013. Tips and tricks about *Shigella* invasion of epithelial cells. Current Opinion in Microbiology 16:32–37.

52. Valencia-Gallardo CM, Carayol N, Tran Van Nhieu G. 2015. Cytoskeletal mechanics during Shigella invasion and dissemination in epithelial cells. Cellular Microbiology 17:174–182.

53. Kühn S, Bergqvist J, Gil M, Valenzuela C, Barrio L, Lebreton S, Zurzolo C, Enninga J. 2020. Actin Assembly around the Shigella-Containing Vacuole Promotes Successful Infection. Cell Reports 31.

54. Shafaei A, Rees J, Christophersen CT, Devine A, Broadhurst D, Boyce MC. 2021. Extraction and quantitative determination of bile acids in feces. Analytica Chimica Acta 1150:338224.

55. Ramos RJ, Zhu C, Joseph DF, Thaker S, LaComb JF, Markarian K, Lee HJ, Petrov JC, Mon-zur F, Buscaglia JM, Chawla A, Small-Harary L, Gathungu G, Morganstern JA, Yang J, Li J, Pamer EG, Robertson CE, Frank DN, Cross JR, Li E. 2022. Metagenomic and Bile Acid Metabolomic Analysis of Fecal Microbiota Transplantation for Recurrent Clostridiodes Difficile and/or Inflammatory Bowel Diseases. Medical Research Archives 10.

56. do Nascimento PGG, Lemos TLG, Almeida MCS, de Souza JMO, Bizerra AMC, Santiago GMP, da Costa JGM, Coutinho HDM. 2015. Lithocholic acid and derivatives: Antibacterial activity. Steroids 104:8–15.

57. Kisthardt SC, Thanissery R, Pike CM, Foley MH, Theriot CM. 2023. The microbial-derived bile acid lithocholate and its epimers inhibit Clostridioides difficile growth and pathogenicity while sparing members of the gut microbiota. Journal of Bacteriology 205:e00180–23.

58. Sorg JA, Sonenshein AL. 2010. Inhibiting the Initiation of Clostridium difficile Spore Germination using Analogs of Chenodeoxycholic Acid, a Bile Acid. Journal of Bacteriology 192:4983–4990.

59. McKenney PT, Yan J, Vaubourgeix J, Becattini S, Lampen N, Motzer A, Larson PJ, Dannaoui D, Fujisawa S, Xavier JB, Pamer EG. 2019. Intestinal Bile Acids Induce a Morphotype Switch in Vancomycin-Resistant Enterococcus that Facilitates Intestinal Colonization. Cell Host & Microbe 25:695–705.e5.

60. Kragh KN, Tolker-Nielsen T, Lichtenberg M. 2023. The non-attached biofilm aggregate. Commun Biol 6:898.

61. Ménard R, Sansonetti PJ, Parsot C. 1993. Nonpolar mutagenesis of the ipa genes defines IpaB, IpaC, and IpaD as effectors of Shigella flexneri entry into epithelial cells. Journal of Bacteriology 175:5899–5906.

62. Ménard R, Sansonetti P, Parsot C. 1994. The secretion of the Shigella flexneri Ipa invasins is activated by epithelial cells and controlled by IpaB and IpaD. The EMBO Journal 13:5293–5302.

63. Sauer K, Stoodley P, Goeres DM, Hall-Stoodley L, Burmølle M, Stewart PS, Bjarnsholt T. 2022. The biofilm life cycle: expanding the conceptual model of biofilm formation. Nat Rev Microbiol 20:608–620.

64. Frick I-M, Mörgelin M, Björck L. 2000. Virulent aggregates of Streptococcus pyogenes are generated by homophilic protein–protein interactions. Molecular Microbiology 37:1232–1247.

65. Pseudomonas aeruginosa interacts with epithelial cells rapidly forming aggregates that are internalized by a Lyn-dependent mechanism - Lepanto - 2011 - Cellular Microbiology - Wiley Online Library. https://onlinelibrary.wiley.com/doi/10.1111/j.1462-5822.2011.01611.x. Retrieved 18 November 2025.

66. Feltham L, Moran J, Goldrick M, Lord E, Spiller DG, Cavet JS, Muldoon M, Roberts IS, Paszek P. 2024. Bacterial aggregation facilitates internalin-mediated invasion of Listeria monocytogenes. Front Cell Infect Microbiol 14.

67. Anderson MC, Vonaesch P, Saffarian A, Marteyn BS, Sansonetti PJ. 2017. *Shigella sonnei* Encodes a Functional T6SS Used for Interbacterial Competition and Niche Occupancy. Cell Host & Microbe 21:769–776.e3.

68. Caboni M, Pédron T, Rossi O, Goulding D, Pickard D, Citiulo F, MacLennan CA, Dougan G, Thomson NR, Saul A, Sansonetti PJ, Gerke C. 2015. An O Antigen Capsule Modulates Bacterial Pathogenesis in Shigella sonnei. PLOS Pathogens 11:e1004749.

69. Datsenko KA, Wanner BL. 2000. One-step inactivation of chromosomal genes in Escherichia coli K-12 using PCR products. Proceedings of the National Academy of Sciences 97:6640–6645.

70. Jaumouillé V, Francetic O, Sansonetti PJ, Tran Van Nhieu G. 2008. Cytoplasmic targeting of IpaC to the bacterial pole directs polar type III secretion in Shigella. The EMBO Journal 27:447–457.

71. Jumper J, Evans R, Pritzel A, Green T, Figurnov M, Ronneberger O, Tunyasuvunakool K, Bates R, Žídek A, Potapenko A, Bridgland A, Meyer C, Kohl SAA, Ballard AJ, Cowie A, Romera-Paredes B, Nikolov S, Jain R, Adler J, Back T, Petersen S, Reiman D, Clancy E, Zielinski M, Steinegger M, Pacholska M, Berghammer T, Bodenstein S, Silver D, Vinyals O, Senior AW, Kavukcuoglu K, Kohli P, Hassabis D. 2021. Highly accurate protein struc-ture prediction with AlphaFold. Nature 596:583–589.

72. Mirdita M, Schütze K, Moriwaki Y, Heo L, Ovchinnikov S, Steinegger M. 2022. ColabFold: making protein folding accessible to all. Nat Methods 19:679–682.

73. Morris, G. M., Huey, R., Lindstrom, W., Sanner, M. F., Belew, R. K., Goodsell, D. S., & Olson, A. J. (2009). AutoDock4 and AutoDockTools4: Automated docking with selective receptor flexibility. Journal of Computational Chemistry, 30(16), 2785–2791.

74. Kim S, Chen J, Cheng T, Gindulyte A, He J, He S, Li Q, Shoemaker BA, Thiessen PA, Yu B, Zaslavsky L, Zhang J, Bolton EE. 2023. PubChem 2023 update. Nucleic Acids Res 51:D1373–D1380.

75. Trott, O., & Olson, A. J. (2010). AutoDock Vina: improving the speed and accuracy of docking with a new scoring function, efficient optimization, and multithreading. Journal of Computational Chemistry, 31(2), 455–461.

76. Humphrey W, Dalke A, Schulten K. 1996. VMD: Visual molecular dynamics. Journal of Molecular Graphics 14:33–38.

