## Supplementary Figures for "Lithocholic acid induces T3SS-dependent formation of invasion-competent *Shigella flexneri* aggregates"

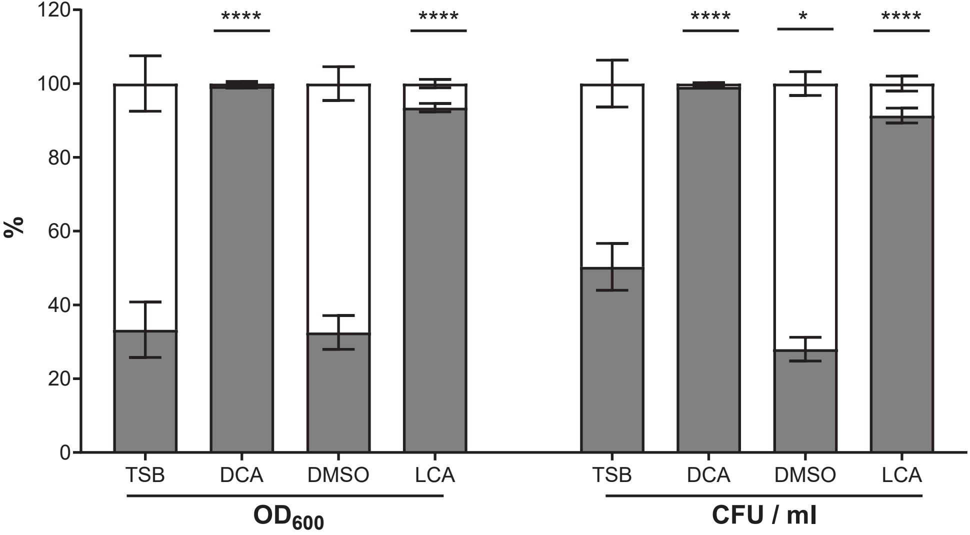


**Figure S1. Comparative quantification of aggregation by $OD_{600}$ and CFUs following the sedimentation assay.** Sedimentation assay of *S. flexneri* in TSB containing DCA (2500 µM), LCA (50 µM), and vehicle for LCA (DMSO). Graph shows mean ± SD from three biological repeats; white, planktonic %; gray, aggregate %. Statistics: one-way ANOVA with Dunnett's multiple comparison; TSB control for DCA; DMSO control for LCA. *, $p < 0.05$; ****, $p < 0.0001$.

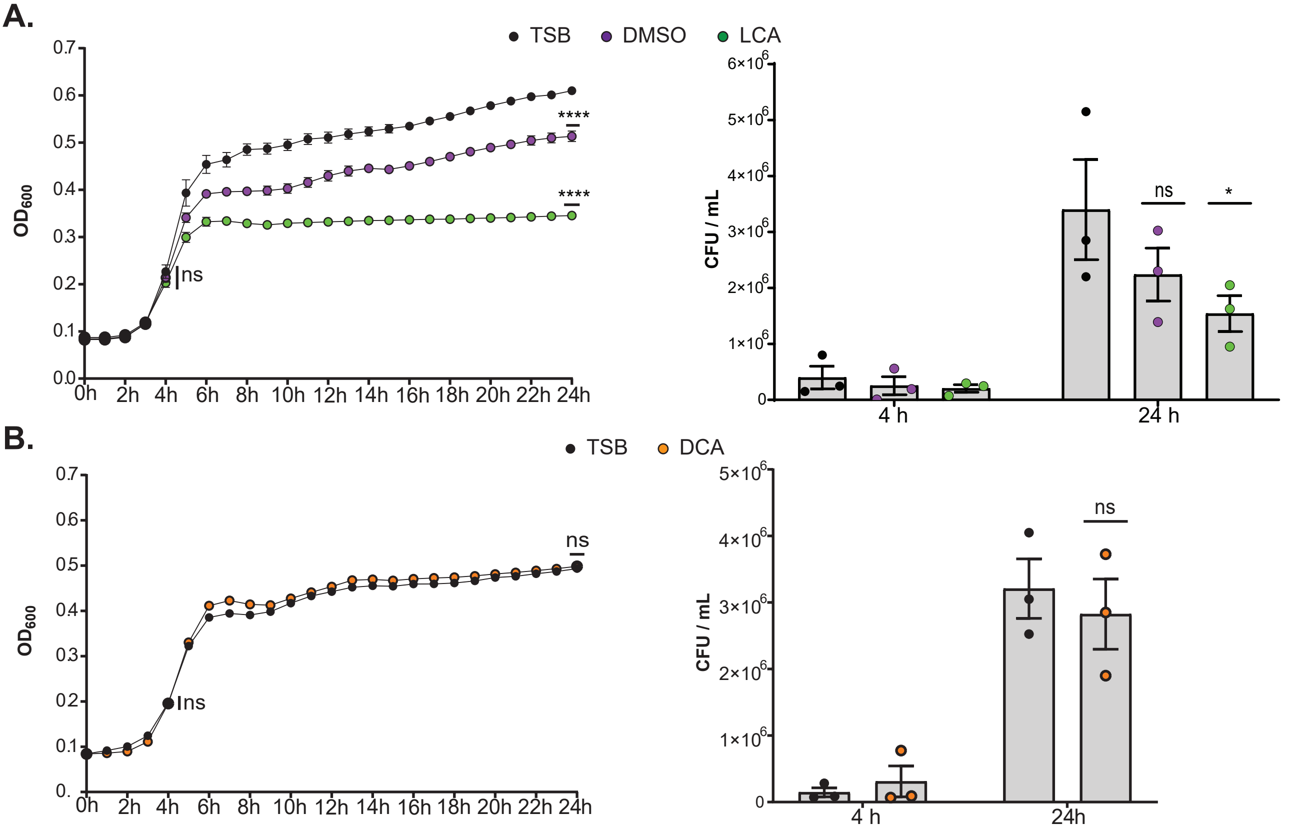


**Figure S2. LCA modestly reduces *S. flexneri* viability at stationary phase.** Left: Growth curves of static *S. flexneri* cultures grown ± 50 μM LCA (A) or ± 50 μM DCA (B) at 37 °C; $OD_{600}$ was measured hourly for 24 h. Right: CFUs were enumerated at 4 h and 24 h to validate $OD_{600}$ measurements. Graphs show mean ± SEM from three biological repeats. Statistics: one-way ANOVA with Dunnett's multiple comparison test relative to the control at each time point. *, $p < 0.05$; ****, $p < 0.0001$; ns, not significant.

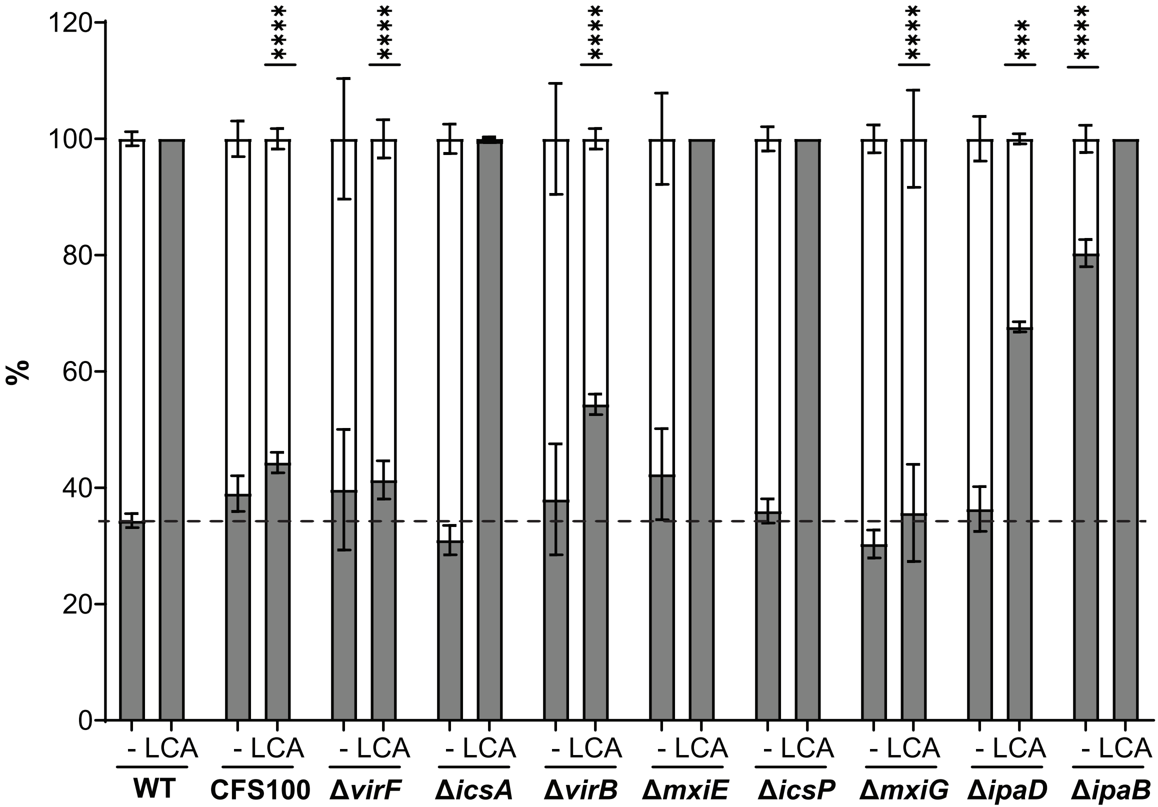


**Figure S3. LCA at 100 μM induces *S. flexneri* aggregation via the T3SS and IpaD.** Sedimentation assay of *S. flexneri* mutant strains grown in TSB supplemented with 100 μM LCA or DMSO vehicle (-). Graph shows mean ± SD from three biological repeats; white, planktonic %; gray, aggregate %. Dashed line indicates basal bile acid-independent aggregation. Statistics: one-way ANOVA with Dunnett's multiple comparison test relative to the corresponding WT control. ***, $p < 0.001$; ****, $p < 0.0001$.

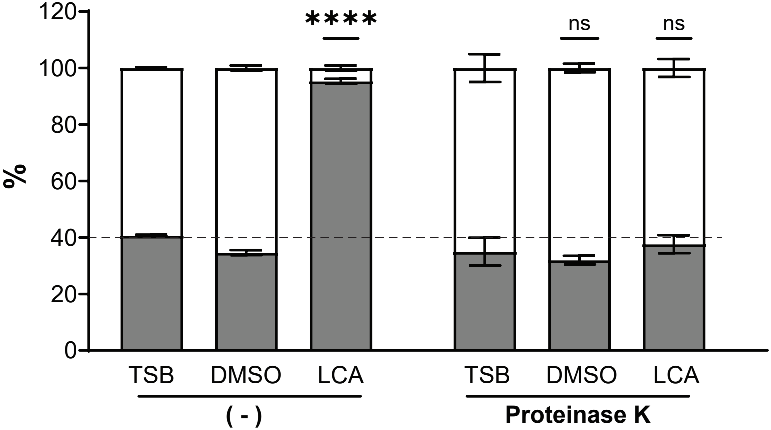


**Figure S4. Proteinase K sensitivity of LCA-induced aggregation.** Sedimentation assay performed in TSB, DMSO, and 50 μM LCA ± proteinase K. Graph shows mean ± SD from three biological repeats; white, planktonic %; gray, aggregate %. Dashed line indicates basal bile acid-independent aggregation. Statistics: one-way ANOVA with Dunnett's multiple comparison test relative to DMSO control. ***, $p < 0.001$; ns, not significant.

A.

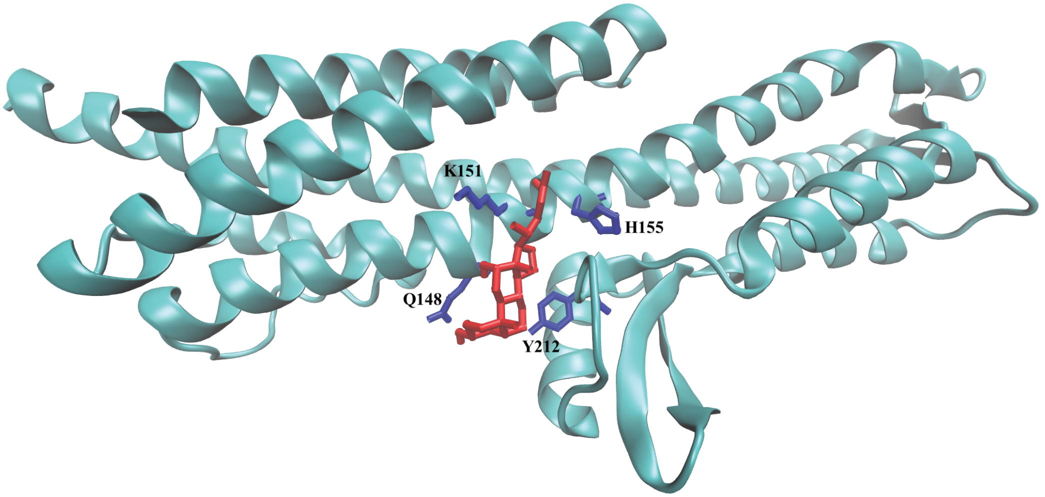


B.

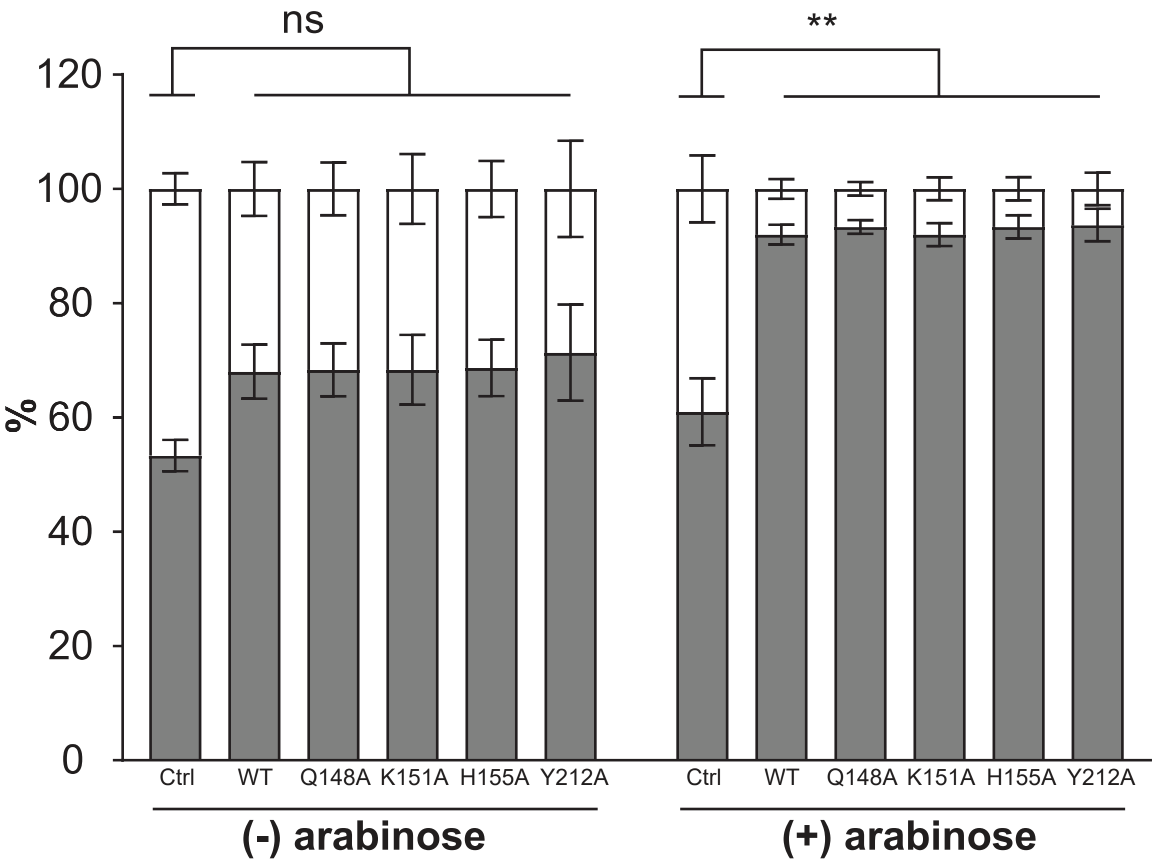


**Figure S5. Potential IpaD residues interacting with LCA.** (A) AlphaFold-predicted structure of IpaD rendered in NewCartoon representation (cyan), with LCA (docked via AutoDock Vina) shown in red bonded representation and predicted ligand-interacting residues Q148, K151, H155 and Y212 highlighted in blue bonded representation. Visualization generated with VMD 1.9.4. (B) Sedimentation assay with Δ*ipaD* mutant harboring pBAD18 (Ctrl), pIpaD-HA (WT), and single alanine substitutions at Q148, K151, H155, and Y212 residues ± 0.2% arabinose. Graph shows mean ± SD from three biological repeats; white, planktonic %; gray, aggregate %. Statistics: one-way ANOVA with Dunnett's multiple comparison test. **, $p < 0.01$; ns, non-significant.
